# Microglial TNFα controls synaptic GABAARs, sleep slow waves and memory consolidation

**DOI:** 10.1101/2022.02.21.481254

**Authors:** Maria Joana Pinto, Lucy Bizien, Julie M.J. Fabre, Nina Ðukanović, Valentin Lepetz, Fiona Henderson, Marine Pujol, Romain W. Sala, Thibault Tarpin, Daniela Popa, Antoine Triller, Clément Léna, Véronique Fabre, Alain Bessis

## Abstract

Microglia sense the changes in their environment. How microglia actively translate these changes into suitable cues to adapt brain physiology is unknown. We reveal an activity-dependent regulation of cortical inhibitory synapses by microglia, driven by purinergic signaling acting on P2RX7 and mediated by microglia-derived TNFα. We demonstrate that sleep induces microglia-dependent synaptic enrichment of GABA_A_Rs in a manner dependent on microglial TNFα and P2RX7. We further show that microglia-specific depletion of TNFα alters slow waves during NREM sleep and blunt memory consolidation in sleep-dependent learning tasks. Together, our results reveal that microglia orchestrate sleep-intrinsic plasticity of synaptic GABA_A_Rs, sculpt sleep slow waves and support memory consolidation.

---

Microglia, the immune cells of the brain, tune neuronal networks in the healthy brain by finely modulating synapses^1^. They can sculpt developing circuits and remodel neuronal connectivity in adulthood by adjusting synapse density, function and plasticity^2–6^. Microglia accomplish these functions either by direct interaction with synaptic elements^5,7^ or through the release of factors^4,8^. Many of the latter, such as prostaglandin, BDNF, IL-1β or TNFα, control synaptic plasticity and were independently shown to regulate sleep^9^. Recent work highlights the ability of microglia to control sleep duration^10,11^; however, whether and how microglia-released factors shape sleep via modulation of synaptic plasticity remains unknown.

Sleep drives plasticity of excitatory synapses. In the cortex, spine turnover during sleep is associated to learning and memory^12,13^ and, in parallel, wake-associated strengthening of excitatory synapses is downscaled during sleep^14,15^. Synaptic inhibition is critically involved in sleep generation and sleep oscillations^16–18^, but whether and how inhibitory synapses are dynamically modulated during sleep remains poorly understood. Here, we uncover the molecular pathway underlying sleep-intrinsic microglia-dependent modulation of synaptic GABA_A_R in light vs. dark periods. Moreover, we demonstrate that microglial TNFα shapes sleep slow waves and supports memory consolidation.

## Daily modulation of GABA_A_Rs in a sleep- and microglia-dependent manner

We first analyzed the modulation of synapses in the frontal cortex in light vs. dark by measuring the synaptic content of neurotransmitter receptors (fig. 1 and fig. S1 and S2). Mice sleep more during the light phase and spend most of the dark phase awake. Therefore, we compared brains of mice at Zeitgeber time 18 (ZT18; dark/middle of wake phase) and at ZT6 (light/middle of sleep phase; fig. 1a). We focused on cortical layer 1 (L1) which is a key node for wide-scale cortical computation^19^. Consistent with the well-established downscaling of excitatory synapses during sleep^14^, synaptic accumulation of the AMPA receptor subunit GluA2 was modestly but significantly decreased at ZT6 as compared to ZT18 (fig. S1). Of note, this level of change is comparable to previous analysis of daily changes in synaptic AMPARs^14,20^. Yet, analysis of L1 inhibitory synapses revealed an increased enrichment of synaptic, but not extra-synaptic, GABA_A_R-γ2 and - α1 subunits at ZT6 (fig. 1a-c and fig. S2b-d), as well as an enhanced proportion of inhibitory synapses containing a GABA_A_R cluster (fig. S2e). No changes were observed in the total signal of GABA_A_Rs (fig S2g), supportive of redistribution of the receptors rather than changes in expression. In contrast, in layer 5 (L5) the synaptic content of GABA_A_Rγ2 did not significantly differ between ZT6 and ZT18 (fig. 1e and fig. S2f). These results reveal a novel form of regulation of GABAergic synapses in light vs. Dark periods with synaptic enrichment of GABA_A_Rs in L1 during the light phase (ZT6), which likely contributes to upregulation of inhibitory transmission in the upper cortex during this phase^21^.

**Fig. 1.**
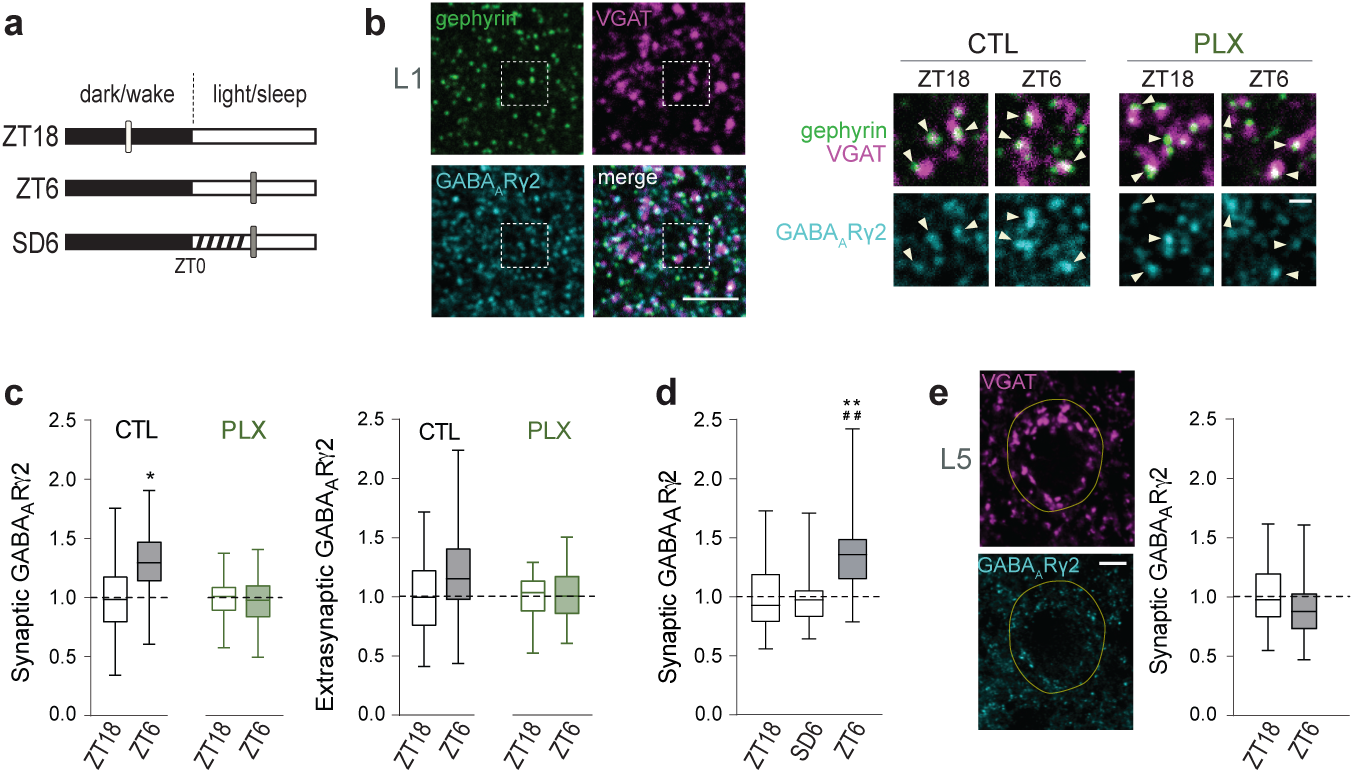
Plasticity of GABA_A_R in light vs. dark is sleep- and microglia-dependent. **a**, Experimental groups: mice at dark phase (ZT18), mice at light phase (ZT6) and mice submitted to sleep deprivation (dashed) in the light phase (SD6). Vertical bars: time of perfusion. **b**, Representative images showing enrichment of GABA_A_Rγ2 (cyan) at cortical L1 inhibitory synapses in the light phase (ZT6). Arrowheads: GABA_A_Rγ2 clusters at gephyrin^+^VGAT^+^ synapses in control (CTL) or PLX3397-treated mice (PLX). Scale bars, 5 and 1 µm. Dashed box corresponds to CTL at ZT18. **c, d,** Mean intensity of GABA_A_Rγ2 clusters at gephyrin^+^VGAT^+^ synapses (synaptic) and at extrasynaptic sites normalized to ZT18. n= 48 to 65 fields of view (FOVs) from 4-5 mice per group. *p<0.05, nested t-test; **p<0.01 compared with ZT18 and ^##^p<0.01 compared with SD6, nested one-way ANOVA followed by Sidak’s multiple comparison test. **e**, Left: Representative confocal images of VGAT and GABA_A_Rγ2 in cortical L5. Yellow line delineates soma identified by NeuN staining. Scale bar, 5 µm. Right: Mean intensity of GABA_A_Rγ2 at somatic VGAT^+^ clusters in L5 normalized to ZT18. n= 60 FOVs from 5 mice per group.

To discriminate between a sleep-intrinsic or time of day-dependent regulation of synaptic GABA_A_R in L1, mice were forced to stay awake during their normal sleep period (sleep deprivation from ZT0 to ZT6, SD6; fig. 1a). The synaptic content of GABA_A_R was not different between mice at ZT18 and mice sleep-deprived in the light phase (SD6; fig. 1d), showing that modulation of synaptic GABA_A_R in L1 is driven by sleep-dependent mechanisms. Sleep deprivation has previously been shown to increase GABA_A_Rs located around excitatory somas^22^. This suggests that the expression of GABA_A_Rs is differentially regulated depending on their subcellular localization.

We have previously shown that microglia control the accumulation of receptors at inhibitory synapses in the spinal cord^23^. This prompted us to investigate the involvement of microglia in the modulation of synaptic GABA_A_R in light vs. dark periods. Strikingly, microglia depletion by feeding mice with the CSF1R antagonist PLX3397^24^ (PLX; fig. S2a, b) completely prevented the changes in synaptic GABA_A_Rs content between ZT6 and ZT18 (fig. 1c and fig. S2b,e). Importantly, as already shown^10,11^ microglia depletion does not alter sleep during the light phase (fig S2h-I; table S2), discarding the possibility that lack of synaptic GABA_A_Rs enrichment upon PLX3397 treatment results from perturbed sleep during the light phase. This shows that the sleep-dependent modulation of synaptic GABA_A_R in L1 requires microglia. Moreover, microglia depletion also reversed the reduction of synaptic AMPA receptor subunit GluA2 at ZT6 (fig S1), which agrees with the recent demonstration that light/dark changes in excitatory synaptic strength are microglia-dependent^10^, a function that has been attributed to CX3CR1 signaling^10^ and that was not further explored in this study.

## Modulation of synaptic GABA_A_R by microglial P2RX7-TNF*α* signaling via CaMKII

We next sought to identify the molecular pathway underlying this novel microglia-dependent synaptic regulation. To accomplish so, we identified the molecular candidates regulating synaptic GABA_A_Rs enrichment in an *ex vivo* paradigm of plasticity (fig. 2) and we subsequently validated these candidates *in vivo* (fig. 3). During sleep, excitatory synaptic plasticity is triggered by NMDAR-dependent dendritic calcium spikes in L1^13^. We thus selected an NMDA-induced GABAergic plasticity protocol, known as inhibitory long-term potentiation (iLTP). This form of plasticity, known to drive potentiation of excitatory synapses^25^, also leads to an upregulation of synaptic GABA_A_Rs^26^ in pyramidal neurons specifically at somatostatin interneurons inputs (SOM-IN)^27^, which are mainly located in L1^28^ (fig. S3). In agreement with the SOM-IN topographical organization, NMDA-induced iLTP in brain organotypic slices led to specific synaptic enrichment of GABA_A_Rs in L1, but not in L5 somatic synapses (fig. 2a, b and fig. S4b-d). This protocol thus mimics *ex-vivo* the synaptic regulations occurring in light vs. dark, and we used it to identify the actors of the L1-restricted sleep-dependent synaptic enrichment of GABA_A_Rs (fig. 1). We further demonstrated that this form of GABA_A_R plasticity was completely abolished when microglia were depleted by PLX (fig. 2a, b and fig. S4a, c). We ruled out possible secondary effects of PLX by showing the same effect upon microglia depletion using Mac1-Saporin (SAP) (fig. S4a) or inactivation using minocycline (fig. 2b).

**Fig. 2.**
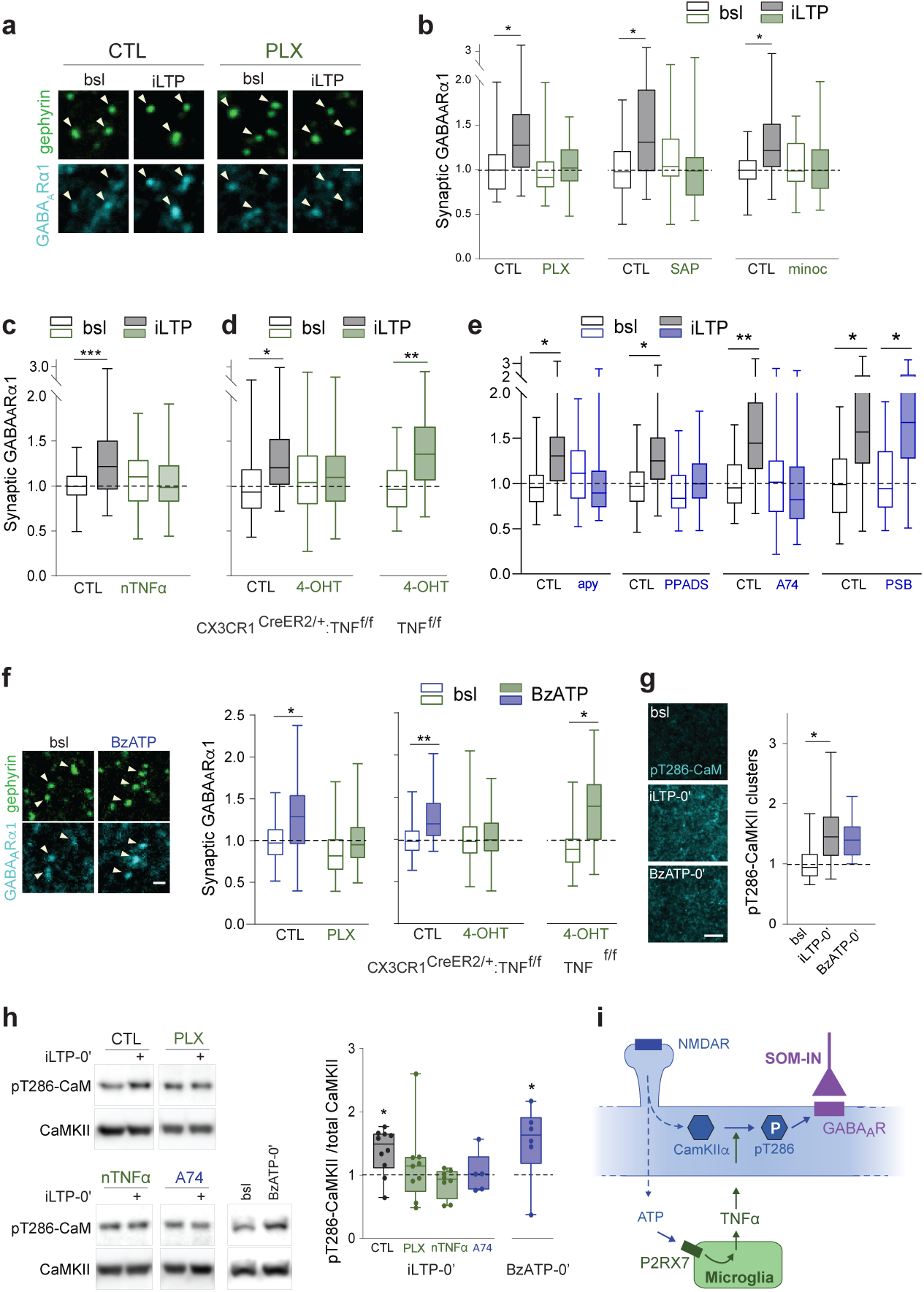
Microglial P2RX7 -TNF*α* signaling drives GABA_A_Rs synaptic enrichment through CaMKII*α* phosphorylation. **a**, Representative confocal images showing increase of GABA_A_Rα1 (cyan) at gephyrin^+^ clusters (arrowheads) upon NMDA-induced inhibitory long-term potentiation (iLTP: 2 min 20 µM NMDA/10 µM CNQX plus 20 min recovery) in organotypic slices cortical L1 (CTL: control; bsl: baseline). **b-e**, Mean intensity of GABA_A_Rα1 clusters at gephyrin^+^ cluster normalized to CTL at bsl. n= FOVs/ independent experiments: (b) n= 44 to 69/ 5 to 6; (c) n= 47 to 53/ 5; (d) n= 66 to 102/ 5 to 7; (e) n= 49 to 68/ 7 to 9. *p<0.05, **p<0.01 and ***p<0.001, nested one-way ANOVA followed by Sidak’s multiple comparison test. iLTP-induced synaptic GABA_A_R enrichment is abolished by: **b**, microglia depletion/inactivation (PLX: PLX3397; SAP: Mac1-saporin; minoc: minocycline); **c**, neutralization of TNFα (nTNFα); **d**, microglia-specific TNFα deletion through 4-hydroxy-tamoxifen (4-OHT)-induced recombination on CX3CR1^CreERT2/+:^TNF^f/f^ but not on a TNF^f/f^ background; **e**, ATP hydrolysis (apy), P2XR antagonist (PPADS) and P2RX7 antagonist (A74), but not P2Y12R inhibitor (PSB). **f**, Left: Confocal images of GABA_A_Rα1 (cyan) at gephyrin^+^ clusters (arrowheads) in bsl and upon BzATP treatment. Right: Mean intensity of GABA_A_Rα1 clusters at gephyrin^+^ clusters normalized to CTL at bsl. n= 49 to 63/ 5 to 6. *p<0.05, **p<0.01, nested one-way ANOVA followed by Sidak’s multiple comparison test. **g**, Left: Thr286-phosphorylated CaMKII is enhanced in L1 at the induction phase of plasticity (iLTP0’ or BzATP0’). Scale bar, 5 µm. Right: Mean intensity of Thr286-phosphorylated CaMKII puncta normalized to bsl. n= 25 to 33 FOVs from 3 independent experiments. *p<0.05, one-way ANOVA followed by Sidak’s multiple comparison test. **h**, Left: Western blot analysis showing iLTP0’-induced CaMKII Thr286-phosphorylation. Right: Ratio between Thr286-phosphorylated CaMKII and total CaMKII normalized to the respective iLTP-free control. n= 5 to 9 independent experiments. *p<0.05 compared with respective control, Kruskal-Wallis test followed by Dunn’s multiple comparisons test. **i**, Model. ATP released downstream NMDA-induced neuronal activity activates microglial P2RX7 with triggers the release of microglial TNFα. TNFα signaling gates CaMKIIα autophosphorylation which controls the enrichment of synaptic GABA_A_Rs in pyramidal neurons.

**Fig. 3.**
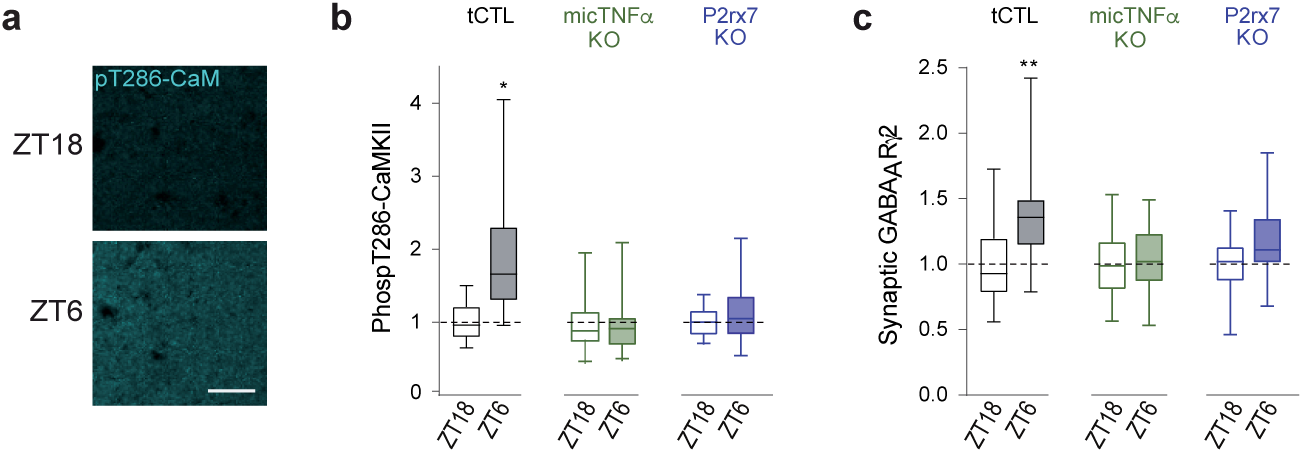
P2RX7 and microglial TNF*α* promote daily changes in synaptic GABA_A_R content and CaMKII phosphorylation. **a**, Representative confocal images of Thr286-phosphorylated CaMKII immunoreactivity in L1 showing higher intensity at ZT6 than ZT18 in transgenic control mice (tCTL). Scale bar, 20 µm. **b**, Mean intensity of Thr286-phosphorylated CaMKII signal in L1 normalized to ZT18 for tCTL, microglia-specific TNFα depletion (micTNFα-KO) and P2rx7-KO mice. n= 37 to 50 FOVs from 4-5 mice per group. **c**, Mean intensity of GABA_A_Rγ2 clusters at gephyrin^+^VGAT^+^ synapses normalized to ZT18. n= 48 to 65 FOVs from 4-5 mice per group. *p<0.05 and **p<0.01, nested t-test.

Microglia produce a broad repertoire of signaling molecules that regulate synaptic function^1^. TNFα, which is mostly if not exclusively produced by microglia in the brain^29^, controls basal synaptic strength^30^, neurotransmitter receptors’ dynamics and homeostatic synaptic plasticity^31^. We thus hypothesized that microglial TNFα controls the activity-dependent regulation of synaptic GABA_A_R. Indeed, neutralization of TNFα by specific antibodies (fig. 2c) as well as conditional microglia-specific TNFα depletion (fig. 2d and fig. S5a) prevented the enrichment of synaptic GABA_A_R upon iLTP. Soluble TNFα derives from the cleavage of a membrane form of TNFα by the TNFα-converting enzyme (TACE)^32^. Both forms can signal through TNFR1 whereas TNFR2 is only activated by membrane TNFα^33^. The increase of synaptic GABA_A_Rs on iLTP-treated slices was prevented by TAPI-1, a TACE inhibitor, and by a TNFR1-but not by a TNFR2-neutralizing antibody (fig. S5b). Finally, we showed that recombinant TNFα and activation of TNFR1 were sufficient to increase synaptic accumulation of GABA_A_Rα1 (fig. S5c-f). These experiments show that microglial soluble TNFα acting through TNFR1 mediates the activity-dependent regulation of GABA_A_R in L1.

We next identified the signaling pathway between neuron and microglia leading to TNFα release upstream of GABA_A_Rs regulation. We first demonstrated that the well-established neuronal CX_3_CL1-microglial CX_3_CR1 signaling, with known roles in synaptic plasticity^34^, is not involved (fig. S4e). Microglia behavior is finely tuned by ATP via an array of purinergic receptors^35^. Microglial cells rapidly react to ATP following the activation of neurons by glutamate and NMDA^36–38^, and the stimulation of microglial P2RX7 promotes the release of TNFα^39^. We thus investigated whether NMDA-induced release of ATP causes microglia-mediated modulation of synaptic GABA_A_R. Indeed, apyrase, a promoter of ATP hydrolysis, PPADS, a broad P2X antagonist, or A740003, a specific P2RX7 antagonist, prevented the increase of synaptic GABA_A_R upon iLTP. In contrast, PSB0739 a specific inhibitor of P2RY12^40^, had no effect on synaptic GABA_A_R upon iLTP suggesting that P2RY12 is not involved in this regulation (fig. 2e). Moreover, BzATP, a P2RX7 agonist^41^ was sufficient to increase postsynaptic GABA_A_Rα1 in L1 (fig. 2f and fig. S6a,b) but not in L5 (fig. S6c). Of note, BzATP had no effect when microglia were depleted or when TNFα was specifically inactivated in microglia (fig. 2f). Thus, ATP/P2RX7 signaling acts upstream of microglial TNFα release to modulate synaptic accumulation of GABA_A_R.

Finally, we explored the intracellular signaling downstream of microglial TNFα. CaMKIIα is a central neuronal mediator of postsynaptic plasticity whose activity is triggered by Ca^2+^/calmodulin and can be prolonged in a Ca^2+^-independent manner by its autophosphorylation at Thr286^42^. Upon iLTP, Thr286-autophosphorylated CaMKIIα leads to insertion of GABA_A_Rs at synapses^43^. We reasoned that TNFα may modulate CaMKIIα Thr286-phosphorylation, as observed in non-neuronal cells^44^. Indeed, Thr286-phosphorylation of CaMKII was increased in L1 upon iLTP (fig. 2g) and this increase was abolished by microglial depletion and by blocking P2RX7 or TNFα signaling (fig. 2h). In agreement, BzATP, which upregulates the synaptic content of GABA_A_Rs via a microglial relay, was sufficient to enhance CaMKII Thr286-phosphorylation (fig. 2g, h). The finding that CaMKII Thr286 phosphorylation, and therefore its activity, is gated by microglial TNFα downstream neuronal activity emphasizes a cardinal position of microglia in fine-tuning synapses.

Collectively, our results support a bidirectional neuron-microglia crosstalk underlying activity-driven GABAergic potentiation in L1 (fig. 2i): ATP released downstream neuronal activity activates microglial P2RX7 with concomitant release of TNFα which modulates CaMKII autophosphorylation and thereby enrichment of synaptic GABA_A_Rs. Noteworthy, our results do not exclude the possible involvement of other cell types acting between neurons and microglia.

## Sleep-dependent modulation of synaptic GABA_A_R driven by P2RX7 and microglial TNFα

Having shown that microglial P2RX7/TNFα pathway controls L1 activity-dependent GABA_A_R plasticity via CaMKII autophosphorylation *ex-vivo*, we examined the involvement of this pathway in the sleep-dependent modulation of synaptic GABA_A_R in light vs. dark (fig. 1). We first showed that CaMKII Thr286-phosphorylation, but not total CaMKII levels, was increased in L1 at ZT6 in a sleep-dependent manner (fig. 3a, b). Notably, such regulation was not found in L5 (fig. S7b). Next, we showed that Thr286-phosphorylation of CaMKII, as well as the synaptic content of GABA_A_Rs in L1, were not enhanced at ZT6 in mice with microglia-specific TNFα depletion (micTNFα-KO, fig. S5a) and in P2RX7-KO mice (fig. 3a-c and fig. S7c, d). Importantly, loss of enhancement at ZT6 is likely not attributed to disturbed sleep since deletion of P2RX7^45^ or deletion of microglial TNFα (as discussed in the next section, figure 4) do not cause major changes in baseline sleep. This shows that P2RX7 and microglial TNFα are required for daily fluctuations of CaMKII Thr286-phosphorylation and synaptic GABA_A_R in cortical L1, which occur in a manner dependent on sleep in the light phase.

**Fig. 4.**
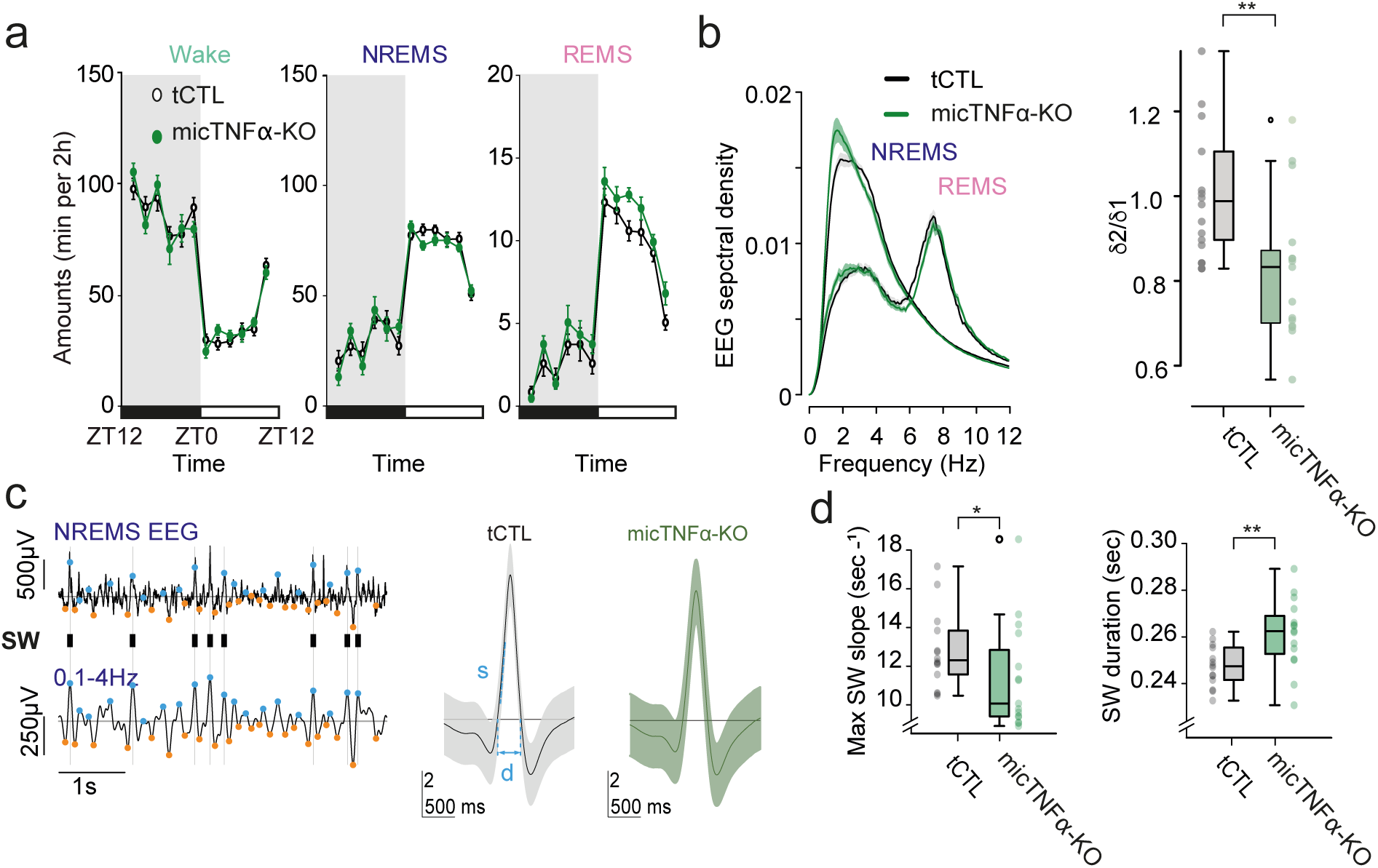
Microglial TNF*α* modulates slow waves during NREMS. **a**, Amounts of vigilance states over 24h reported by 2h segments. Wake, NREMS and REMS are not significantly different between tCTL and micTNFα-KO mice (n=15 mice per group; two-way RM-ANOVA, p=0.1490; p=0.2784, p=0.6838 respectively) **b**, Left: Average spectral density (top) of tCTL and micTNFα-KO (lines: means. Shaded area: SEM) and distribution of peak frequency of delta oscillations in a 24h time period. Right: Ratio between power in faster delta frequencies (δ2, waves from 2.5 to 3.5 Hz) and slower delta frequencies (δ1, waves from 0.75 to 1.75 Hz) in NREM sleep (Mann-Whitney W = 169, p = 0.0043). **c**, Left: Examples of EEG and 0.1-4Hz filtered EEG during NREMS from a transgenic control tCTL; the positive and negative peaks of the delta-filtered signal are indicated by orange and blue points, respectively. Ticks on grey lines indicate large positive deflections corresponding to Slow Waves (SW). Right: grand average SW for tCTL(black) showing duration (d) and maximum slope (s), and micTNFα-KO (green). **d**, Characteristics of SW in a 24h time period. micTNFα-KO mice exhibit significantly shorter peak onset slope and longer SW duration than tCTL mice. Mann-Whitney W=42 p=0.005 and W=153, p=0.37 respectively. n = 14 tCTL and 15 micTNFα-KO respectively

## NREM slow waves shaped by microglial TNF*α*

The results above show that microglial TNFα is required for a novel sleep-dependent regulation of inhibitory synapses in cortical L1. Sleep is an alternation of REM and NREM sleep periods which are hallmarked by major EEG oscillatory patterns^46^. During NREM sleep, slow waves in the delta frequency band (0.1-4Hz), quantified as slow wave activity (SWA), result from the synchronous alternation of active (up) and silent (down) states of cortical neurons. Cortical GABAergic inhibition is a major actor of NREM sleep slow waves^47,48^. In particular, the activation of somatostatin positive interneurons, that mainly target L1, triggers down-states^17,18^. Because microglial TNFα modulates synaptic GABA_A_R at L1, we anticipated that it could also shape SWA. Sleep was analyzed using EMG and epidural EEG recordings, and we concentrated our analysis in the frontal cortex where slow waves are predominant^49^. We first showed that microglial TNFα has limited effects on sleep-wake patterns as shown by the lack of major alterations in the amounts of wake, NREM and REM sleep between micTNFα-KO and tCTL mice along a light/dark cycle (fig. 4a).

We then analysed the spectral density of the EEG during sleep (figure 4b). In agreement with a role of microglial TNFα in the regulation of SWA, we found a shift in the EEG spectral density towards lower frequency activity during NREM sleep with a slower peak frequency in the delta range in micTNFα-KO as compared to tCTL (p=0.02539), with no change in the delta power (Wilcoxon test: W = 67, p=0.06) (fig. 4b). Faster frequencies in the delta range (δ2, waves from 2.5 to 3.5 Hz) during NREM sleep, but not lower frequencies (δ1, waves from 0.75 to 1.75 Hz), reflect prior sleep-wake history^50^. By computing the ratio of δ2 over δ1 power, a clear decrease is observed in micTNFα-KO (fig. 4b right), further supportive of a redistribution of SWA to a slower frequency range. There were no detectable differences in the other frequency ranges nor in REM sleep (fig. 4b). Finally, we explored how microglial TNFα shapes the properties of individual slow waves during NREM sleep. Slow waves were identified in the epidural EEG as alternation of large transient negative and positive deflections in the 0.1-4 Hz filtered EEG (fig. 4c). We verified that these deflections correspond to up and down states respectively (fig. S8; Nir et al. 2011).

Remarkably, the maximum ascending slope of the slow waves, which coincides with the onset of the cortical positive deflection (downstate; fig. S8c), was decreased (∼17 % decrease in micTNFα-KO as compared to tCTL, fig. 4d left) and the duration of the SW down state (fig. S8a, b) was increased in mice depleted of microglial TNFα as compared to controls (fig. 4d). These results indicate that microglial TNFα shapes slow waves during NREM sleep by favouring transition into the down states.

## Microglial TNF*α* tunes memory consolidation

Slow waves are known to play a causal role for the consolidation of memory during sleep^51,52^. Our previous results show that microglial TNFα exquisitely shapes slow waves during NREM sleep, leading us to anticipate a crucial role of microglial TNFα in the memory consolidation processes during sleep. To test this hypothesis, we compared the consolidation of different learning tasks known to be sleep-dependent^53–55^ in micTNFα-KO and tCTL mice. In a first learning paradigm^53^, mice learnt to run on top of a complex wheel attached to an accelerating rotarod in a first 20-trial session (S1). After one day of ad libitum sleep, performance on the complex wheel was assessed on a second 20-trial session by measuring the latency to fall (S2; fig. 5a, b).

**Fig. 5.**
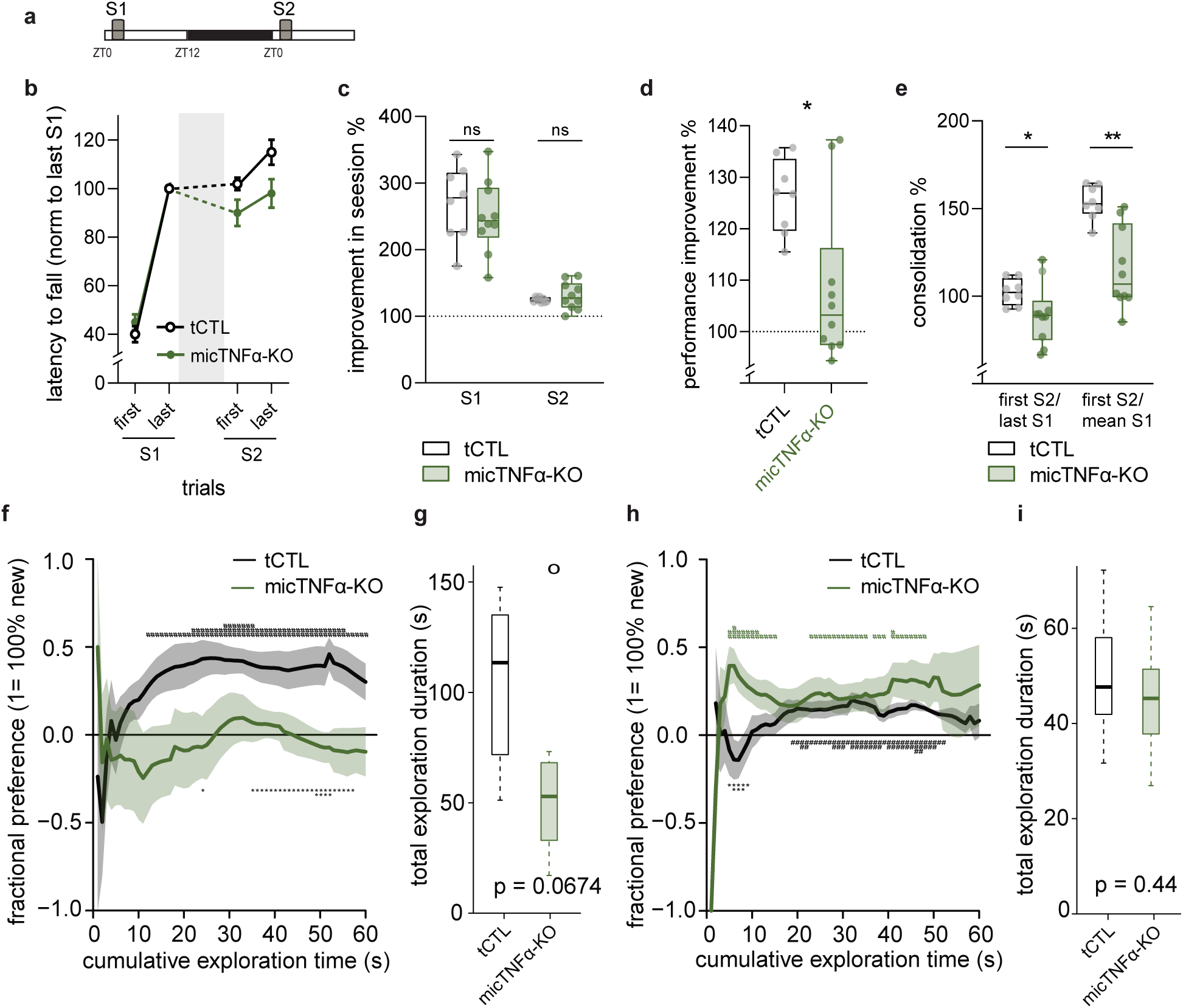
Microglial TNF*α* required for memory consolidation in sleep-dependent learning tasks. **a**, Experimental design: mice learn to run on the complex wheel (session 1, S1) and consolidation of memory is tested the following day (session 2, S2). Between S1 and S2, mice are left undisturbed in their cages. **b**, Average latency to fall off the complex wheel in the first three and the last three trials of S1 and S2 from tCTL and micTNFα-KO. Grey area represents undisturbed sleep-wake cycle. Dashed line represents S1 to S2 consolidation. **c**, Improvement within each session measured as the ratio between mean of the best three trials and first three trials. **d**, Performance improvement across sessions measured as the ratio between mean of S2 and S1 trials. ***p<0.001, Mann-Whitney test. **e**, Consolidation of motor learning across sessions measured in two ways: ratio between mean of first 3 trials of S2 and mean of last 3 trials of S1 (first S2/ last S1) or ratio between mean of first 3 trials of S2 and mean of S1 trials (first S2/ mean S1). *p<0.05 and **p<0.01, Mann-Whitney test. (b-e) 8 tCTL and 10 micTNFα-KO mice. **f, h,** Novelty preference in the novel floor-texture recognition (FTR) (f) or the Novel Object Recognition (NOR) task (h). Fractional preference is expressed as a function of cumulative time of exploration (see Methods). The data represent the mean+/- sem of the preference for the object placed on the novel floor texture (f) or for the novel object (h) computed for each animal. ^#^p<0.05, ^##^p<0.01 and ^###^p<0.001 preference for one object vs. no preference and *p<0.05, **p<0.01 and ***p<0.001 CTL vs. micTNFα-KO. Unpaired t tests. The fractional preference is equal to 1 or -1 when the animal only explored respectively the object placed on the novel or familiar floor texture in the FTR and the novel or familiar object in the NOR, and 0 when the animal spent exactly the same amount of time on the two objects. **g, i,** Total duration of exploration of the novel floor (FTR; g) and of the novel object (NOR; i). Boxes represent quartiles and whiskers correspond to the range of data; points are singled as outliers if they deviate more than 1.5 x inter-quartile range from the nearest quartile. g, °p= 0,0674 CTL vs. micTNFα-KO, Unpaired t tests. **i,** p= 0,44 CTL vs. micTNFα-KO. Unpaired t tests; (f-i) n= 11 tCTL and 8 (f,g) or 16 (h,i) micTNFα-KO mice.

We first confirmed that the learning, evaluated as the improvement of performance within each session, was not different between tCTL and micTNFα-KO (fig. 5c). This shows that lack of microglial TNFα did not alter complex motor learning, which is known to be sleep-independent^53^. We further showed that it does not impair locomotor activity and does not induce anxiety-like behavior as assessed in an open-field task (fig. S9). We next measured the improvement of performance <u>in the complex motor learning task</u> between S1 and S2 and found that it was higher in tCTL as compared with micTNFα-KO (fig. 5d). Finally, we tested the memory consolidation by comparing performance at the beginning of the second session (First S2) with either the last or the mean performance of the first session (Last S1 or Mean S1). In agreement with our initial hypothesis, memory consolidation was impaired in mice lacking microglial TNFα as compared to controls (fig. 5e). Remarkably, in this complex motor learning task, the improvement of performance between sessions and the memory consolidation are both known to be sleep-dependent processes^53^. To confirm the involvement of microglial TNFα in memory consolidation, tCTL and micTNFα-KO mice performed a floor-texture recognition task, which is known to depend on cortical activity during NREM sleep^54^. Mice first explored an arena with a single floor-texture (training session, smooth or rough) and containing two identical objects on each side. After one day of ad libitum sleep, mice explored the same arena containing the same two objects which were placed either on a familiar or on a novel floor-texture (testing session, smooth and rough). As expected^54^, tCTL mice spent more time exploring the object located on the novel floor-texture during the testing session (fig 5f). In contrast, micTNFα-KO mice explored significantly less the object located on the novel floor-texture (fig 5f,g) confirming the impairment of memory consolidation on these mice.

Finally, tCTL and micTNFα-KO mice were tested in the novel-object recognition task (fig 5h, i). During the sampling session, mice explored two identical objects, and the next day (testing session) one of the original objects is replaced by a novel object. The amount of time taken to explore the novel object during the testing session provides a quantification of memory consolidation. Noteworthy, however, both genotypes exhibited larger amount of time spent exploring the novel object (fig 5i) indicating normal consolidation for this test.

## Discussion

Microglia, the principal immune cells of the brain, are now acknowledged as instrumental for the perception of the external environment^56^. How microglia actively translate their external sensing into suitable cues to adapt circuitries in the healthy brain is yet unknown. Our results favour a model in which microglia sense neuronal activity through an ATP/P2RX7 signalling pathway and respond to it by releasing TNFα. Microglial TNFα then gates the phosphorylation of neuronal CaMKII that modulates GABA_A_R content at layer 1 cortical synapses (fig. 2). In the light period, microglia act through TNFα to upregulate synaptic GABA_A_R content in L1 (figs. 1 and 3), which likely contributes to the strengthening of inhibitory transmission in the upper cortex^21^. In line with a prominent role of inhibition in the generation of slow waves^17,18,47,48,57^ microglial TNFα shapes slow waves by controlling transition into down states (fig. 4). We finally demonstrate that microglial TNFα is determinant for memory consolidation previously shown to occur in a sleep-dependent manner^53,54^ (fig. 5).

TNFα has long been known as a sleep factor. Administration of exogeneous TNFα promotes SWA and NREM sleep^58,59^ whereas inhibition of endogenous TNFα reduces NREM sleep^refs^ ^in^ ^60^. Similarly, intracerebral or intraperitoneal injection of TNFα^61^ supresses REM sleep, which is increased in a dark period-specific manner in TNFα KO mice^62^. These results seem to partially contradict our results showing no difference in NREM sleep amount and no dark-specific increased REM sleep in mice with microglia-specific TNFα depletion. However TNFα injection either directly in the brain^59^ or intraperitoneally^58^ leads to concentrations that are likely higher than the physiological concentration^63^, with putative widespread diffusion and off-target and indirect effects. Furthermore, the studies using TNFα or TNFα receptors knock-out have not used cell-specific and inducible inactivation and it is thus difficult to discriminate between microglial specific effect to indirect effects due to non-microglial TNFα and/or secondary developmental alterations. Finally, TNFα is also a major mediator of inflammation, which induces sleep dysfunction by yet elusive mechanisms^64^. Our work now provides possible molecular and cellular links between sleep and inflammation.

Our results were obtained from experiments performed both *in vivo* and in organotypic slices. The latter system closely mimics the *in vivo* brain by preserving tissue architecture and cellular composition. In these slices microglia retain their 3D ramified morphology and their functional properties^65–67^. They further conserve their ability to regulate synapses^23,68^. This indicates that functional interactions between neurons and microglia are conserved in organotypic slices. Indeed, in this work, we have identified microglial TNFα, ATP/P2RX7 and CaMKII as molecular actors of synaptic GABA_A_Rs regulation in organotypic slices and we have further confirmed their role *in vivo* in the light vs. dark regulation of GABA_A_Rs plasticity.

Excitatory synapses are scaled down during sleep, through removal of AMPA receptors, to compensate for potentiation due to ongoing learning during wake^69^. Microglia may contribute to downscaling during sleep by eliminating excitatory synapses^70^, a behaviour presumably tuned down during wake by noradrenaline^71,72^. More recently, microglial CX3CR1 signalling was shown to differentially regulate excitatory synaptic transmission along the light/dark cycle^10^. We now show that microglia tune a sleep-dependent regulation of inhibitory synapses restricted to cortical layer 1, via mobilization of the P2RX7/TNFα signalling and regulation of synaptic GABA_A_R content. Noteworthy, comparable amplitude of changes in the GABA_A_R was found in the sleep/wake cycle analysis of the proteome of forebrain synaptosomes^20^. The layer 1 vs layer 5 specificity may arise from the molecular difference of GABAergic synapses across the somato-dendritic arbour as proposed^27^. Alternatively but not exclusively, it may result from a differential expression of TNF-R1 along the cortical layers and/or from layer-specific behavior of microglia^73^. Altogether, microglia are arising as active players in excitatory and inhibitory synapse remodelling along the sleep/wake cycle, likely to underlie daily oscillations in synaptic strength^15,21,74^.

Our study suggests that microglia can target specific sets of synapses by acting locally in a sleep-dependent manner; however, the mechanisms supporting this spatiotemporal specificity have remained elusive. Microglia expression of TNFα mRNA peaks at the middle of the light phase^75^, when sleep is maximal. Specificity could be achieved by differential microglial production of TNFα; however, our attempt to visualize distribution of TNFα protein in the cortex at sleep and wake states failed due to unreliability of anti-TNFα antibodies (not shown). Dynamics of TNFα release by microglia along a light/dark cycle in specific sleep states thus awaits the advent of suitable tools. Another potential source of specificity may come from sleep-specific patterns of neuronal activity and downstream ATP release that can then engage local microglia. For instance, intense calcium spikes occur during sleep on the apical dendrites of pyramidal neurons inducing plasticity in upper cortical layers^13^. ATP levels in frontal cortex actually peak in the light period in a sleep-dependent manner^76^. Resolving the mechanism promoting spatiotemporal specificity of microglia functions during sleep would give great insight into microglia-neuronal crosstalk in the physiological brain.

Cortical inhibition is involved in the control of slow wave activity during NREM sleep^17,57,77^, with a prominent role in the onset of down states^48,78^. Such control is likely achieved through inhibitory networks in the upper layers of the cortex. Thalamic drive onto L1 inhibitory neurogliaform cells induces transition into down states^47^. In addition, SOM-IN densely target L1 and their firing precedes entry into down states^18^. Accordingly SOM-IN chemoactivation increases by about 17% the slope of slow-waves and triggers down-states^17^. In agreement with the ability of microglial TNFα to control inhibitory synapses in L1, we found that depletion of microglial TNFα leads to a 17% decrease in slow-waves slope. Remarkably, the slope of slow waves is under homeostatic control. Indeed, sleep deprivation increases the slope of slow-waves^17,50^, and in the early sleep, when sleep need is high, the slope of the slow-waves is steeper by 15% above the average^79^. Therefore, the decrease in slow wave slope in micTNFα-KO mice aligns with our earlier finding that these mice exhibit reduced expression of sleep need^80^. In further support, lack of microglial TNFα alters SWA toward slower frequencies with a reduction in the range of SWA frequencies (δ2) shown to be related to sleep homeostasis^78^.

In this study we show that L1 inhibitory synapses are modulated by microglia in a sleep-dependent manner. Our data favor the possibility that the microglia-targeted GABAergic synapses arise from SOM-IN. However, inhibition on L1 also originates from local axonal arbors of neurogliaform and canopy cells^19^ and the identity of the presynaptic neurons remains to be established. In agreement with their ability to modulate inhibitory synapses in cortical L1, yet without ruling out their possible involvement in other brain regions and/or by other mechanisms, microglia are regulators of slow waves during NREM sleep through TNFα. Together with evidence of astrocytic involvement in slow wave activity^81,82^, our work offers insight into how local glial modulation of neuronal networks may tune brain oscillations. <u>L</u>ocal modifications in SWA within cortical regions involved in learning processes are crucial for learning consolidation and performance improvement<u>^83,84^</u>. <u>O</u>ur work suggests that microglia could play a role in facilitating the necessary adjustments in SWA during NREM sleep that accompany the intense recruitment of neural networks during learning episodes.

Cortical inhibition controls sleep-dependent memory consolidation^77^, which itself critically depends on slow waves^51,52^. It has even been proposed that one of the major physiological role of sleep is to allow memory consolidation^85^. We now show that mice lacking microglial TNFα display impaired memory consolidation when tested in a complex rotarod motor learning task or in the floor-texture recognition (FTR) task. In these two tasks, memory consolidation is sleep-dependent^53,54^. This suggests a possible involvement of microglial TNFα in the sleep processes that promote memory consolidation, while leaving open the possibility of a non-sleep-dependent mechanism. Strikingly, memory consolidation was not impaired in micTNFα-KO in the novel object recognition (NOR) task. These phenotypes mimicked that of mice lacking nNOS specifically in SOM-IN that also display altered performance in FTR but not in NOR^77^. This congruence strengthens the hypothesis that microglial TNFα is a major actor of the regulation of cortical inhibition underlying consolidation of memories during sleep.

Finally, this work adds to the yet limited knowledge of microglia functions in the healthy adult brain^4,6^ and establishes microglia as genuine regulators of inhibitory synapses, brain oscillations and memory in the healthy brain. Noteworthy, microglial regulation likely occurs through the control of CaMKIIα, which is a cardinal regulator of synaptic plasticity^42^. It may occur on SOM-IN inputs, which are involved in the control of complex behaviours such as sleep, but also decision making and learning^86^. Finally, this work demonstrates that microglia tune slow waves and support memory consolidation probably by acting during sleep^85^. We thus anticipate a far wider involvement of microglia in other forms of plasticity and higher brain functions.

## Acknowledgments

This work was funded by: Agence nationale pour la recherche: SYNTRACK -R17096DJ (AT); Agence nationale pour la recherche: EXPECT 17-CE37-0022-2 (CL, DP); European Research Council: PLASTINHIB Project No: 322821 and MICROCOPS (AT); Human Brain Project: HBP SGA2-OPE-2018-0017 (AT); EMBO fellowship: ALTF 362-2017 (MJP);

We gratefully acknowledge Astou Tangara, Benjamin Mathieu and the IBENS imaging facility (IMACHEM-IBiSA), member of the French National Research Infrastructure France-BioImaging (ANR-10-INBS-04), which received support from the "Fédération pour la Recherche sur le Cerveau - Rotary International France" (2011) and from the program « Investissements d’Avenir » ANR-10-LABX-54 MEMOLIFE.

We are grateful to Amandine Delecourt, Eleonore Touzalin, Deborah Souchet for excellent technical assistance.

We thank Sonia Garel (IBENS, Paris), Etienne Audinat (Institut de génomique fonctionnelle, Montpellier) and Lauriane Ulmann and François Rassendren (Institut de génomique fonctionnelle, Montpellier) for providing CX_3_CR1^CreERT2^ and Ai9; TNF^flox^ and P2rx7 -KOs respectively.

We are grateful to the IBPS Animal facility for breeding and care, the IBPS-NPS Phenotypic core facility, and particularly Jean Vincent.

## Author contributions

Conceptualization: AB, AT, CL, DP, MJP, VF. Formal Analysis: CL, MJP, VF. Funding acquisition: AB, AT, CL, DP, VF. Investigation: JF, FH, LB, MJP, NÐ, VF, AB, MP, RWS, TT Software: CL, VL, MJP. Writing – original draft: AB, MJP. Writing – review & editing: AB, AT, CL, DP, MJP, VF, RWS.

Authors declare that they have no competing interests.

