## Supplemental information and figures for "Microglial TNFα controls synaptic GABAARs, sleep slow waves and memory consolidation"

**This PDF file includes:**

Materials and Methods

Figs. S1 to S9

Tables S1-S2

**Other Supplementary Materials for this manuscript include the following:**

None

### Materials and Methods

#### Animals and housing

Mice were housed under standard conditions (12 h light/dark cycle; lights on at 7:00 A.M.). All experiments were performed in conformity with the European Committee Council Directive 86/609/EEC and were approved by the local Charles Darwin Ethical Committee (Ce5-2014-001; 1339-2015073113467359 and 2018022121466547). $CX_3CR1^{GFP}$(*1*); $CX_3CR1^{CreERT2}$ (*2*); $TNF^{flox}$ (*3*); P2X7R-KO (*4*); $SOM^{iCre}$ (*5*) and Ai9 (*6*) mouse lines were housed at the animal facility of Institut de Biologie de l'ENS or at the animal facility of Institut Biologie Paris Seine (Paris, France). $CX_3CR1^{GFP}$, $CX_3CR1^{CreERT2}$ and Ai9 were kindly provided by Sonia Garel, $TNF^{flox}$ by Etienne Audinat and P2X7R-KOs by François Rassendren. Constitutive TNFα knockout mice (TNFα-KO) were generated by crossing $TNF^{flox}$ males with PGK1-Cre females (*7*). C57BL/6J pregnant females and wild-type males were obtained from Janvier Labs.

#### Brain organotypic slices

Brain organotypic slices were prepared as previously described (*8*). Brains were removed from P3–P7 C57BL/6J mice (wild-type, $CX_3CR1^{+/+}$:$TNF^{f/f}$ or $CX_3CR1^{CreERT2/+}$:$TNF^{f/f}$) and hemispheres separated in dissection medium (33 mM glucose in PBS). Coronal slices (350 µm) were cut using a McIllwain tissue chopper (Mickle Laboratory), and placed on Millicell cell culture inserts (Millipore, PICM03050) (2 to 3 slices per insert). Slices were maintained at 37°C in 5% $CO_2$/air for 14 to 21 d in MEM (GIBCO, 21090-022) supplemented with 20% heat-inactivated horse serum (GIBCO, 16050-122), 2 mM glutamine, 10 mM glucose, 20 mM HEPES, 10 U/mL penicillin and 10µg/mL streptomycin. Culture medium was changed 3 times per week.

#### Adult primary microglia cultures

Isolation of primary microglia from adult mouse brain was performed as previously described (*9*). In brief, adult mice were anesthetized by inhalation of isoflurane and transcardially perfused with 10 ml PBS. Brains were individually dissociated in solution containing 30-40U papain (Worthington, LK003176), 7.2 U dispase II (Sigma, D4693) and 10 mg/ml DNAse (Sigma, DN25), followed by mechanical dissociation. Microglia were isolated by collecting the 70-37% interphase of a Percoll gradient (Sigma, P4937) and plated on poly-ornithine-coated plates in DMEM/F12 (GIBCO, 31331-028) supplemented with 10% heat-inactivated fetal bovine serum (GIBCO, 10500-064), 10 U/mL penicillin and 10µg/mL streptomycin.

#### Spleen cultures

Adult mice were anesthetized by inhalation of isoflurane and transcardially perfused with 10 ml PBS, followed by dissection of the spleen and its dissociation in 150mM $NH_4Cl$, 10 mM $NaHCO_3$, 0.1 mM EDTA solution.

Spleen cells were kept in culture in DMEM (GIBCO, 61965-026) supplemented with 10% heat-inactivated fetal bovine serum (GIBCO, 10500-064), 1 mM sodium pyruvate, 10 U/mL penicillin and 10µg/mL streptomycin.

**Treatments**

Table 1 - Drugs and neutralizing/activating antibodies used.

| Drug | Concentration used | Source (catalog no.) |
|---|---|---|
| 4-hydroxytamoxifen (4-OHT) | 2 µM | Sigma (H7904) |
| A740003 (A74) | 10 µM | TOCRIS (3701) |
| Apyrase (apy) | 10 U/ml | Sigma (A6535) |
| Armenian Hamster IgG isotype ctr (IgG) | 5 µg/ml | BioLegend (400912) |
| BzATP | 100 µM | Sigma (B6396) |
| CNQX | 10 µM | TOCRIS (1045) |
| KN62 | 5 µM | Hello Bio (HB0359) |
| LPS | 10 µg/ml | Invivogen (tlrl-3pelps) |
| Mac-1-Saporin (SAP) | 1 µM | Advanced Targeting Systems (IT-06) |
| minocycline | 50 nM | Sigma (M9511) |
| NMDA | 20 µM | Sigma (M3262) |
| CX3CL1 neutralizing antibody (n-CX3) | 2 µg/ml | R&D systems (AF472) |
| TNFR1 neutralizing antibody (n-N1) | 5 µg/ml | R&D systems (MAB430) |
| TNFR2 neutralizing antibody (n-N2) | 5 µg/ml | BioLegend (113302) |
| TNFR1 activating antibody (TNFR1A) | 5 µg/ml | R&D systems (AF-425-PB) |
| TNFR2 activating antibody (TNFR2A) | 5 µg/ml | R&D systems (AF-426-PB) |
| TNFα neutralizing antibody (nTNFα) | 1 µg/ml | R&D systems (MAB4101) |
| PLX3397 (PLX) | 10 µM | MedKoo Biosciences (206178) |
| PPADS | 200 µM | TOCRIS (0625) |
| PSB0739 | 1 µM | TOCRIS (3983) |
| Recombinant mouse TNFα | 60 nM | R&D systems (410-MT) |
| TAPI-1 | 50 µM | Millipore (579051) |

Treatments to brain organotypic slices in culture were performed as follows. For long-term treatment with PLX3397 (7 to 9 days), Mac1-saporin (7 days) and 4-hydroxytamoxifen (12 to 15 days), the drug or vehicle were added to the culture medium at each medium change. All other treatments were performed between days *in vitro* (DIV) 16 to 20 in pre-warmed artificial cerebrospinal fluid (aCSF; 125 mM NaCl, 2.5 mM KCl, 2 mM $CaCl_2$, 1 mM $MgCl_2$, 5 mM HEPES, 33 mM glucose, pH 7.3). To guarantee rapid penetration of the drugs into the tissue, slices were immersed in aCSF during treatments (final volume of 2 ml, 1 ml under and 1 ml inside the insert). Induction of iLTP was accomplished by 2 min treatment with NMDA and CNQX followed by 20 min recovery in NMDA/CNQX-free aCSF. BzATP treatment followed the same temporal pattern as iLTP. Whenever indicated, drugs were added before [pre-incubation of 15 min (for A740003, apyrase, fluoroacetate, KN62, minocycline, neutralizing and activating antibodies, PPADS and PSB0739) or 40 min (TAPI-1)] and during iLTP or BzATP treatment. For drugs diluted in DMSO a 1000x concentrated stock was used, and an equal volume of DMSO added to the controls. To remove contaminating $K^+$ from apyrase (*10*), dialysis was done in 0.025 µm pore nitrocellulose membranes (Millipore, VSWP01300).

For immunohistochemistry purposes, brain organotypic slices were fixed in 2% PFA (Electron Microscopy Sciences, 15710-S) in PBS for 1h at 4˚C and washed 3 times with PBS. For immunostainings of synaptic markers, organotypic slices were cryopreserved on 20% sucrose for at least 24h at 4˚C and cut on a cryostat to

14 μm-thick sections. Following the normal flattening of organotypic slices in culture, slices are approximately 150-200 μm thick at the time of fixation, allowing us to cut between 9 to 12 cryostat 14 μm-thick sections per each organotypic slice.

**Analysis of sleep-wake synaptic plasticity**

Sleep-wake experiments were carried out in 12- to 15-week-old C57BL/6J males kept on a 12h:12h light/dark cycle. Wild-type mice were fed with normal or PLX3397-containing food (290 mg PLX3397 in 1 kg standard chow, formulated by SSNIFF, S9555-P710) for 2 weeks. Microglia-TNFα depleted mice (micTNFα-KO, $CX3CR1^{CreERT2/+}$:$TNF^{f/f}$) and transgenic controls (tCTL, $CX3CR1^{GFP/+}$:$TNF^{f/f}$) were fed with tamoxifen-containing food (1000mg tamoxifen citrate in 1 kg of chow, formulated by SSNIFF, A115-T71000) for 6 days. Sleep-wake experiments were performed 2 weeks after tamoxifen feeding in order to allow repopulation of short-lived peripheral $CX3CR1^{+}$ cells thus restricting recombination to microglia (*2*, *11*). Sleep (S) and wake (W) mice were kept undisturbed in their cages and taken at the middle of the light period and at the middle of the dark period, respectively. Enforced wake was accomplished by gentle introduction of novel objects inside the cage and gentle tapping of the cage for 6h starting at the onset of the light period (*12*). At the indicated times, mice were collected and rapidly anesthetized by inhalation of isoflurane, followed by transcardial perfusion with 2% PFA. Brains were dissected, post-fixed at RT for 6h in 2% PFA (Electron Microscopy Sciences, 15710-S) and cryopreserved on 20% sucrose for 5 days at 4˚C. Brains were rapidly frozen and sagittal sections of the left hemisphere were cut on a cryostat at 9 μm thickness for further immunostaining. To guarantee a good sampling, 6 sagital sections (distancing at least 100 μm in the brain tissue) were collected per brain onto the same slide and used for imaging.

**Immunohistochemistry (IHC)**

Cryostat sections were submitted to blocking and permeabilization in 0.25% gelatin-0.1% triton and incubated with primary antibodies at 4˚C overnight with shaking. Slices were then incubated with secondary antibodies at RT for 3h with shaking. Slides were mounted on Vectashield mounting medium (Vector Laboratories, H-1000) containing DAPI (Invitrogen, D3571).

Whenever indicated (see table 2), decloaking chamber-based heat-induced epitope retrieval (HIER) was performed before immunostaining. Sections were processed in a decloaking chamber (Biocare Medical, model DC2008INTL) at 110˚C for 20 min in antigen decloaker solution (Biocare Medical, CB910M), incubated at RT for 15 min with 15% methanol 0.3% $H_2O_2$ in PBS, and incubated at RT for 40 min in 1% sodium borohybride in PBS (solution prepared fresh between 45min to 2h before use) with shaking. Slices were then immunostained as previously described.

Table 2 - Antibodies and IHC method used.

| Antibody | IHC in organotypic slices | IHC in adult brain | Source (catalog no.) |
|---|---|---|---|
| Gephyrin | indirect IHC (1:400) | HIER (1:400) | SySy (147-011) |
| VGAT | indirect IHC (1:500) | HIER (1:500) | SySy (131-004) |
| $GABA_ARα1$ | indirect IHC (1:2500) | HIER (1:500) | SySy (224-203) |
| $GABA_ARγ2$ | indirect IHC (1:500) | HIER (1:500) | SySy (224-003) |
| Iba1 | indirect IHC (1:400) | | Wako (019- 19741) |
| GluA2 | | HIER (1:400) | Millipore (MAB397) |
| Homer1 | | HIER (1:500) | SySy (160-003) |
| NeuN | indirect IHC (1:250) | HIER (1:250) | Millipore (MAB377) |
| CaMKII | | indirect IHC (1:500) | Abcam (ab22609) |
| pT286-CaMKII | | HIER (1:100) | Abcam (ab171095) |

**Confocal imaging and quantitative analysis**

For imaging of synaptic clusters in organotypic slices, a Leica SP8 inverted confocal microscope equipped with a 63x glycerol objective (1.4 NA), a hybrid detector and the Leica Application Suite X (LAS X) software was used. Single confocal plane images were taken to cortical L1 or putative cortical L5 with a 3x zoom factor applied (512 x 512 pixels, pixel size of 120 nm). Images were taken to 3 to 5 cryostat sections obtained from 2 to 3 organotypic slices for each condition of an independent experiment. For imaging of synaptic clusters in adult brain tissue, a Leica SP5 upright confocal microscope equipped with a 63x oil objective (1.4 NA), a hybrid detector and the Leica Application Suite X (LAS X) software was used. Images taken were 1 µm z-stacks of 3 confocal planes (*z* resolution 0.5 µm) with a 4x (at L1) or 5x (at L5) zoom factor applied. Images were taken to the frontal cortex (between 0.5 to 2 mm lateral to midline and > 2 mm anterior to bregma) from 6 cryostat sections for each brain. All imaging was performed on a bidirectional manner at 200 Hz speed.

Double or triple colocalization analysis of synaptic clusters was performed automatically on ICY software (*13*) by a custom-built protocol. Analysis was done on either images of a single confocal plane (organotypic slices) or sum projections of z-stacks (adult brain). First, clusters in each channel were detected by the Spot Detector block (*14*) which extracts spots based on an undecimated wavelet transform detector (size of spots set to scales 2 or 3; threshold values kept constant for each experiment). Identification of colocalization events between detected clusters was accomplished by the object-based colocalization Studio block using the Statistical Object Distance Analysis (SODA) software. In brief, a fixed distance (set to between 500 to 600 nm) from the center of mass of each detected cluster is used to detect colocalization events with clusters from another channel (*15*). For each channel, detected clusters were then pooled into colocalized and non-colocalized and their total number and individual intensity values extracted. In triple-colocalization, a cluster is considered as colocalized when colocalization was detected with the 2 other channels individually. For instance, a $GABA_AR$ cluster is considered synaptic when colocalized with gephyrin clusters or both gephyrin and VGAT clusters for double or triple colocalization, respectively. An $GABA_AR$ cluster is considered extrasynaptic when it shows no colocalization with any of the synaptic markers used. To reduce variability due to uneven background intensity, local background subtraction was applied to each individual cluster.

**ELISA for TNFα**

Following 4-hydroxytamoxifen treatment, $CX_3CR1^{+/+}$:$TNF^{f/f}$ or $CX_3CR1^{CreERT2/+}$:$TNF^{f/f}$ organotypic slices were treated with LPS for 3h in aCSF, lysed in cell extraction buffer [10 mM Tris, 100 mM NaCl, 1 mM EDTA, 1% triton-X100, 10% glycerol, 0.1% SDS supplemented fresh with 1 mM NaF, 2 Mm $Na_3VO_4$ and protease inhibitor cocktail (Roche, 05056489001)] by mechanical dissociation, centrifuged for 10 min at 16000g at 4˚C and the supernatant collected. Adult primary microglia were plated in 96-well plates (40.000 cells per well) and stimulated with LPS or BzATP for 6h. Spleen cells were plated in 24-well plates (3.000.000 cells per well), allowed to deposit for 2h and stimulated with LPS for 24h. For both microglia and spleen cells, medium was collected, centrifuged for 5 min at 10000g at 4˚C and the supernatant collected. Levels of TNFα in the samples were determined by the mouse TNFα uncoated ELISA kit (Invitrogen, 88-7324).

**Western blot**

For western blot analysis of CaMKII phosphorylation, organotypic slices were homogenized in cell extraction buffer by mechanical dissociation, centrifuged for 10 min at 16000g at 4˚C and the supernatant collected. Total protein concentration was determined by the Pierce BCA protein assay kit (ThermoFisher, 23225). Samples (40µg of protein) were denatured at 95˚C for 5 min in denaturating buffer (125 mM Tris-HCl, 10% glycerol, 2% SDS, bromophenol blue, and 5% β-mercaptoethanol added fresh), resolved by SDS-PAGE in 8% polyacrylamide gel and transferred to PVDF membranes. Membranes were blocked in 5% dry milk and incubated with primary antibody against phosphoT286-CamKII (1:1000, Abcam, ab171095) at 4˚C overnight. Membranes were incubated with HRP-conjugated secondary antibody at RT for 1h. Proteins were visualized by chemiluminescence using SuperSignal West Femto Maximum Sensitivity substrate (Thermo Scientific, 34095) and scanned on a ImageQuant LAS 4000 imaging system (GE Heathcare). Membranes were stripped with 0.2 M NaOH for 40min, reprobed for total CamKII (1:1000, Abcam, ab22609) and resolved with Lumi-Light Western Blotting substrate (Roche, 12015196001). Quantification was performed on Image J.

**Electrode implantations for sleep monitoring**

Mice were anaesthetized with ketamine/xylazine before being fixed in a stereotaxic apparatus. All coordinates were adapted from the mouse brain atlas (*16*).

Mice were implanted with the classical set of electrodes (made of enameled nichrome wire, 150 µm in diameter) for polygraphic sleep monitoring (*17*). Briefly, two electrodes were positioned epidurally in holes perforated into the skull: one electrode was inserted over the frontal cortex (2 mm lateral to midline and 2 mm anterior to bregma; left hemisphere), and one electrode was inserted over the cerebellum. Two electrodes were inserted into the neck muscles. This configuration provided a frontal electro-encephalogram (EEG) derivation and an electromyogram (EMG) derivation. All electrodes were soldered to a miniconnector (Antelec, La Queue en Brie, France). Dental acrylic cement was used to anchor the electrodes to the skull. Mice were allowed to recover for 14 days before starting the administration of tamoxifen-containing food for 6 days or PLX3397-containing food for 7 days. Mice were placed in individual recording chambers and connected to the recording system with a light-weight cable and a swivel allowing free movements. Recordings started 2 weeks after tamoxifen administration.

**Sleep recording and analysis**

A 24-hour baseline (BL) recording starting at dark onset was done to analyze sleep-wake stages in 1) $CX3CR1^{CreER2}$:$TNF^{f/f}$ (micTNFα-KO) and $CX3CR1^{GFP/+}$:$TNF^{f/f}$ (control) mice and 2) wild-type mice fed with normal or PLX3397-containing food for 2 weeks. EEG and EMG signals were amplified (2000x), filtered, digitized at 2000 Hz and subsequently down-sampled to 200 Hz by an Embla Module acquisition hardware and the Somnologica acquisition software (Medcare, Reykjavik, Iceland). Polysomnographic recordings were visually scored offline for consecutive 10-s epochs as wake, non REM sleep (NREMS) or REM sleep (REMS) as previously described (Henderson et al., 2016). Briefly, wake was defined by low-amplitude/high frequency EEG activity and elevated EMG tone. NREMS was defined by high-amplitude/low-frequency (<4 Hz) EEG activity and low EMG tone. REMS was characterized by an EEG activity with theta oscillations and a complete absence of EMG tone with occasional twitches. Bouts were defined as consecutive 10-s epochs of similar vigilance state and could be as short as one epoch. Vigilance states amounts for each animal were expressed as minutes per 2- or 12-hr intervals or 24-hr. Sleep architecture was assessed by calculating the bout mean duration and number for each vigilance state.

**Spectral analysis**

A spectrogram was first computed using the modulus of the Fast Fourier Transform of 512 EEG samples at 200 Hz (bouts of 2.56 s) multiplied by a Hanning window; the spectral density for each vigilance state was taken as the median of the spectrogram for all EEG bouts belonging to the vigilance state.

**Slow-waves analysis**

Slow waves were analyzed along the 24h time period. The EEG was first band-pass filtered using a 0-phase Butterworth filter (0.1-4Hz, order 2). The slow waves (SW) were defined as NREMS events accompanied with a large and slow positive deflection of the EEG measured against a reference placed over the cerebellum. We first identified as slow positive peaks of the band-pass filtered signal with the 0-crossings surrounding the peaks of the filtered signal separated by more than 0.4 s and less than 2 s. Amongst these peaks, the SW were selected as the positive oscillations in NREMS larger than a threshold corresponding to: $m+3s$, where $m$ is the median of the positive peaks of the filtered EEG during Wake and $s$ is the difference between the quantiles 0.5 and 0.84 of these peaks; $m$ and $s$ coincide with the mean and the standard deviation if the peak amplitude values would

be normally distributed but this threshold is less sensitive to occasional recording artefacts in the EEG during Wake. The relevance of this definition to track events associated with cortical down-states is confirmed in supplementary figure 8.
The average SW traces were obtained by averaging the EEG centered on the peaks of filtered EEG; the duration of the positive peaks was derived from the time delay between the downward and upward 0-crossing of the filtered EEG around the peaks.

**Linear trans-cortical electrodes**

To confirm the correspondence between the SW as detected by the procedure above with cortical down-states, we implanted 64 channel linear electrodes (Cambridge Neurotech ASSY-156-H3, 64 channels interspaced by 20μm) in the motor cortex of two C57Bl/6J mice. The operations were carried out under isoflurane anaesthesia with the aid of an operating microscope. The body temperature was maintained constant during the surgery (36°C) using a heating pad controlled by a rectal thermometer. General analgesia was assured by subcutaneous injection of buprenorphine (3mg/kg), whereas local analgesia of the skin and skull was provided via subcutaneous injection of lidocaine (0.5 ml of 2 mg/ml solution). Lidocain spray (10%) was used for the ears, before setting the mouse in the stereotaxic frame (Kopf Instruments). After the povidone-iodine (100 mg/ml) and ethanol (70%) disinfection of the surgical area, a sagittal incision of the scalp was performed. Following exposure of the skull, burr holes were drilled above the studied structures. All coordinates (medio-lateral and antero-posterior) were taken relative to bregma, while the depth was measured starting from the dura mater. The electrode was lowered perpendicular to the cortical surface at the coordinates AP: 1.1mm, ML :-0.9mm, z: -1.28mm and the surface covered with Dura-Gel (Cambridge NeuroTech), the rest of the implant is cemented to with dental cement onto the skull after having applied a thin layer of superbond on it. A miniature stainless steel screw was implanted over the cerebellum, serving as electrical reference and ground. The skin ridges were sutured around the connector and mice were allowed to recover in their home cage for at least 7 days. The implanted mice were housed individually in standard cages. Mice were recorded with the Intan RHD USB interface board as recording setup (at sampling frequency 30kHz) in a round openfield (40 x 40 cm) where the nest of the home cage was transferred during 2 hours at noon.

**Linear electrodes processing**

The signal from the linear electrodes was decomposed offline in LFP using a forward and reverse pass filtering with a low-pass filter (0.1-150Hz Butterworth filter of order 2) and in multi-unit activity. The SWs were identified using the same procedure as for EEG applied to the LFP signal from the surface channel of the linear electrodes. The multi-unit activity was extracted offline by first performing a forward and reverse pass of filtering with high-pass filter (500Hz-8kHz, Butterworth filter of order 2), and second extracting the multi-unit spikes selected as the transient negative events deviating more than 6 times the standard deviation of the high-pass filtered signal. The peak of the SW LFP traces was detected from the delta-filtered signal.

**Open-field test**

Mice were placed at the center of an open-field arena (50 × 50 cm) and were allowed to explore it for 15 min. The open-field was divided into a central area (17 x 17 cm) and a peripheral area. Mice were video-tracked with Ethovision software 14.0 by Noldus® (Wageningen, Netherlands) and the distance moved, velocity and time spent in the central area were analyzed for 15 min.

**Complex wheel task**

Sleep-dependent consolidation of motor learning was assessed by the complex wheel task as previously described (*18*). A complex wheel with missing rungs (bar pattern as in (*18*)) was mounted on a rotarod. tCTL and micTNFα-KO mice with no habituation or pre-training on the rotarod were used. Two learning sessions (S1 and S2) were done 24h apart at ZT1, the beginning of the light period. S1 and S2 consisted of 20 and 19 trials on the complex wheel with an acceleration increasing from 0 to 40 rpm in 10 min. Mice were allowed a 5 min rest in their cages between each 10 trials. For each trial, latency to fall corresponds to the time the mouse kept running on the complex wheel without falling. For each mouse, latency to fall off the complex wheel in each S1 and S2 trials was normalized to the average of the last 3 trials in S1. The normalized values were used to plot the graphs in figure 5. Considering that memory consolidation of mice with poorer performances on the complex wheel is less impacted by sleep deprivation than skilled learners (*18*), only the best learners within each group were taken into consideration for the analysis. Within each group (tCTL and micTNFα-KO), best learners correspond to the mice that on S1 performed better than the median of the average performance in all trials. The experimenter was blind to the genotypes.

**Memory recognition tasks**

Sleep-dependent consolidation of recognition memory (*19*, *20*) was assessed by the novel floor-texture recognition (FTR) and novel object recognition (NOR) tasks. FTR and NOR tasks were performed in a square open-field (42cm) and started at ZT3. Both tasks are based on innate novelty preference in mice. Habituation consisted of 1) a 30-min exploration period of the open-field with cagemates (Day1), 2) a 15-min period during which each mouse was placed individually in the empty open-field (Day2), and 3) a 15-min period during which the mice were placed in the open-field with two identical objects (Day3). The tests were executed in the following order: the FTR task was performed on days 4 and 5 and the NOR test on days 6 and 7. On the FTR training session (Day4), mice were allowed to explore two new identical objects in the arena containing a single floor texture (smooth or rough plexiglas) for a 5-min period. This was repeated 2 times (3x5-min exploration period in total). On the testing session (Day5), the mice were tested for the FTR paradigm (5 min) during which the arena contains the same two objects but two floor-textures (smooth and rough) on opposing halves of the floor. On the NOR training session (Day6), mice were allowed to explore two new identical objects for a 5-min period. This was repeated 2 times (3x5-min exploration period in total). On the testing session (Day7), the mice were tested for the NOR paradigm (5 min) during which one of the original objects was replaced with a new object. At the end of each session, mice were replaced and remained in their home cage for 24 hr. Mice that did not accumulate 10 seconds of total inspection time per object over two training sessions in the FRT or the NOR tests were excluded.

Since novelty preference may take some time to become visible (e.g. if the animal takes some time to begin the exploration of one of the two options available) but shall decay along extended exploration (when the novel situation becomes familiar), we chose to examine the object preference as a function of the cumulative time of exploration in the session. For this purpose, we first assessed the amount of time spent exploring each object per 10-sec epochs using the Ethovision software 14.0. Then, we computed the object preference as a function of cumulative time of exploration. For each animal, $v_{nov}(t_i)$ and $v_{fam}(t_i)$ correspond to the amount of time the mouse, at the time in the session $t_i$, spent exploring respectively the object placed on the novel or familiar floor texture in the FTR and the novel or familiar object in the NOR. For each epoch, we compute the cumulative time of exploration $T_j$, $T_j = \sum_{t_i<t_0^j} v_{nov}(t_i) + v_{fam}(t_i)$ and the object preference $p(T_j)$ as $p(T_j) = \left(\sum_{t_i<t_0^j} v_{nov}(t_i) - v_{fam}(t_i)\right)/T_j$ which value shall be 1 or -1 if the animal only explored respectively the object placed on the novel or familiar floor texture in the FTR and the novel or familiar object in the NOR, and 0 if the animal spent exactly the same amount of time on the two objects. The object preference values are linearly interpolated by step of 1 second (of cumulated exploration time) to compute average/sem across animals.

**Statistical analysis**

Statistical significance was defined as follows: ns, not significant (p>0.05); ***p<0.001; **p<0.01 and *p<0.05.

*$GABA_AR$ plasticity:* Results are presented as minimum to maximum box-plots. Graphs and statistical analysis were performed in Prism 8.0 (GraphPad software). Statistical significance was assessed by nonparametric tests. For comparisons of changes between two groups, Nested t-test or Kolmogorov Smirnoff test (for cumulative distributions) were performed. For comparisons between multiple groups, nested one-way ANOVA followed by Sidak's multiple comparison test or Kruskal-Wallis test followed by Dunn's multiple comparisons test were used.

*Sleep recordings:* Sleep data were analyzed using Prism 9.3 (GraphPad Software). Normality and homoscedasticity assumptions were verified prior to the use of any parametric tests (D'Agostino-Pearson normality test or Shapiro-Wilk test and equality of variances *F-test)*. Data violating normality were log transform. Statistics were performed with repeated measures two-way ANOVAs to test the significance between genotypes (tCTL and micTNFα-KO) or treatment (normal or PLX3397-containing food) and time (repeated measures over 2-hr segments) for wake, REMS and NREMS. When appropriate, ANOVAs were followed by the Sidak's multiple comparisons test between genotypes. Two-tailed unpaired t-test or Mann-Whitney test were also employed to assess the significance of the effects of the genotype or treatment on the different variables (amounts, bout mean durations, numbers of bouts) per 12- or 24-hr.

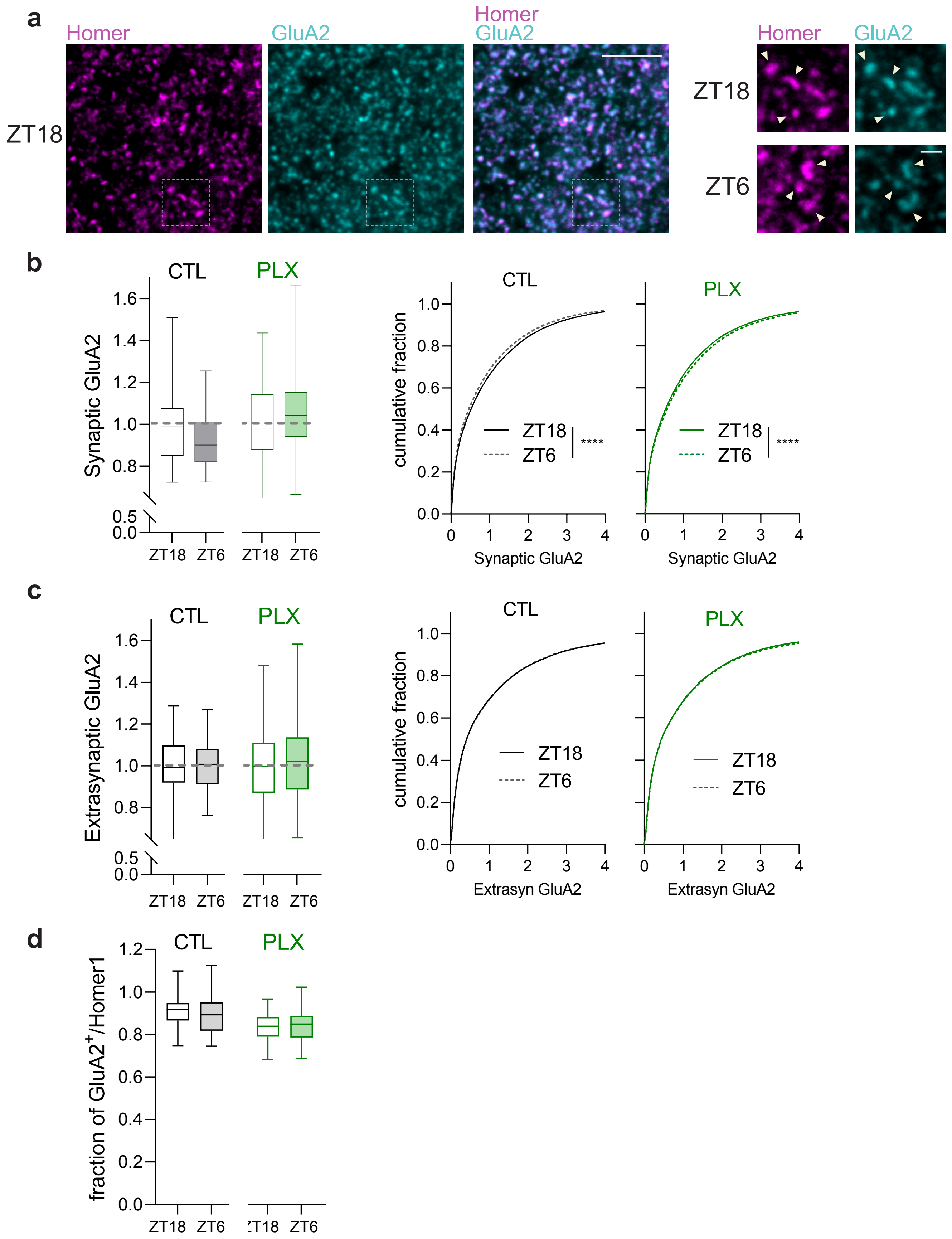


**Figure S1. AMPARs plasticity across the light/dark cycle. a**, Confocal images showing downregulation of GluA2 (cyan) at Homer1⁺ cluster (magenta, arrowhead) at ZT6 in cortical L1. Scale bars, 5 and 1 µm. **b**, Left: Mean intensity of GluA2 clusters at Homer1⁺ clusters. Right: Cumulative fraction of GluA2 clusters' intensities at Homer1⁺ clusters in controls CTL (black) and PLX3397 treated (green) mice. ****p<0.0001, Kolmogorov-Smirnoff test. **c**, Left: Mean intensity of GluA2 at extrasynaptic sites. Right: Cumulative fraction of GluA2 clusters' intensities at extrasynaptic sites in controls CTL (black) and PLX3397 treated (green) mice. **d**, Fraction of Homer1⁺ clusters colocalized with GluA2 in controls CTL (black) and PLX3397 treated (green) mice. b-d**,** n= 46-58 FOVs from 4-5 mice per group.

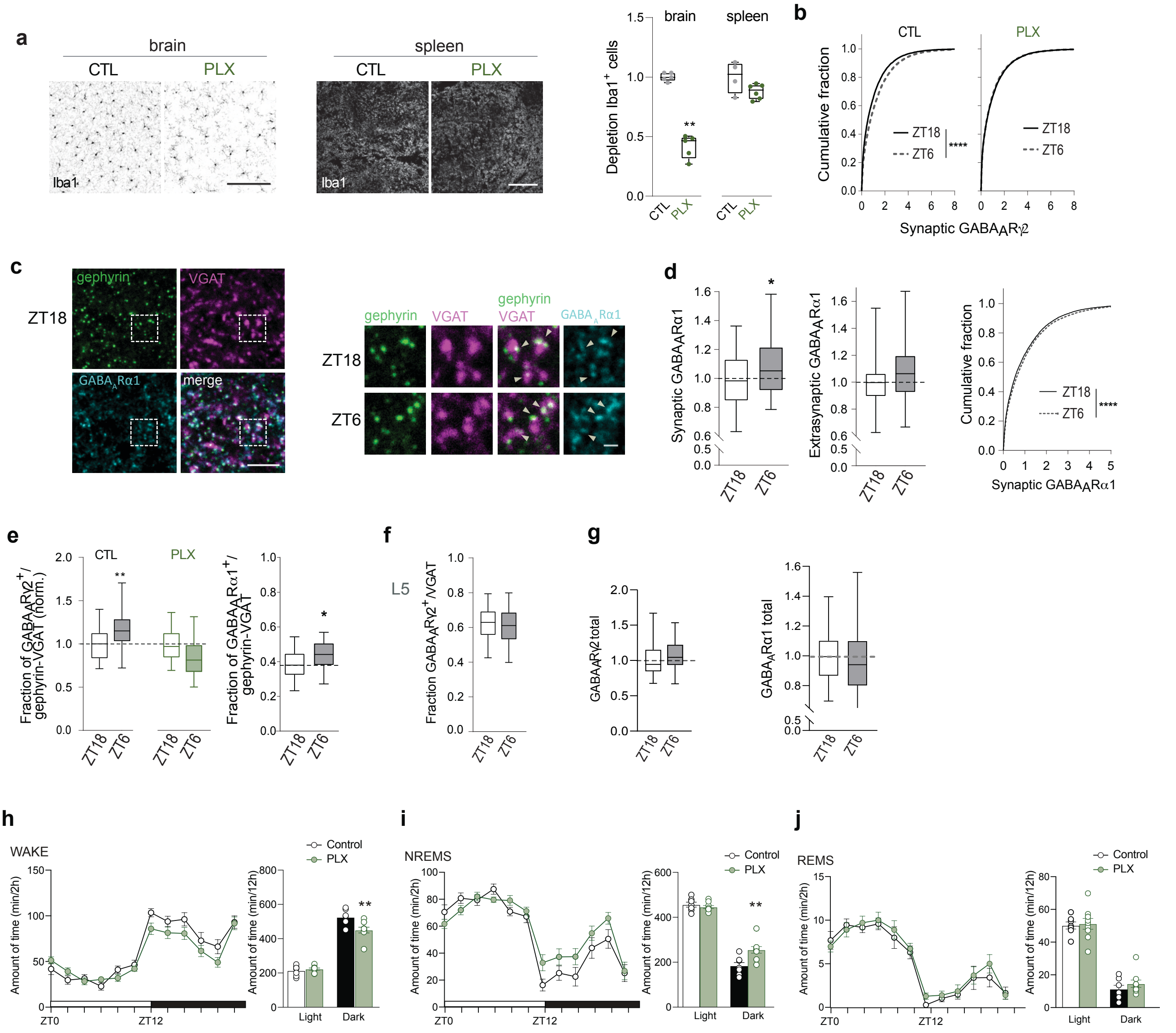


**Figure S2. GABA$_A$Rs plasticity across the light/dark cycle. a**, Left: Confocal images of Iba1$^+$ cells revealing partial depletion in the brain, but not in the spleen, by 2-week feeding with PLX3397-containing food (PLX). Scale bar, 200 µm. Right: Density of Iba1$^+$ cells normalized to CTL. n= 5 mice for CTL and PLX. **p<0.01, Mann Whitney test. Data are mean ± SEM. **b**, Cumulative fraction of GABA$_A$Rγ2 clusters' intensities at gephyrin$^+$VGAT$^+$ synapses in cortical L1. ****p<0.0001, Kolmogorov-Smirnoff. **c**, Representative confocal images showing increase of GABA$_A$Rα1 (cyan) at cortical L1 inhibitory synapses (arrowhead: gephyrin$^+$VGAT$^+$) at ZT6 in comparison to ZT18. Scale bars, 5 and 1 µm. Dashed box corresponds to enlarged detail in ZT18. **d**, Left: Mean intensity of GABA$_A$Rα1 clusters at gephyrin$^+$VGAT$^+$ synapses (synaptic) and extrasynaptic sites. *p<0.05, nested t-test. Right: Cumulative fraction of GABA$_A$Rα1 clusters' intensities at gephyrin$^+$VGAT$^+$ synapses. ****p<0.0001, Kolmogorov-Smirnoff test. **e**, Fraction of gephyrin$^+$VGAT$^+$ synapses colocalized to GABA$_A$Rs clusters. *p<0.05 and **p<0.01, nested t-test. **f**, Fraction of somatic VGAT$^+$ cluster colocalized to GABA$_A$Rγ2 clusters in cortical L5. **g**, Mean intensity of total GABA$_A$Rγ2 (left) and GABA$_A$Rγ2 signal (right). **h-i**, Amounts of vigilance states over 24 hr in controls CTL (black) and PLX3397 treated (green) mice in Wake (h), NREMS (i) and REMS (j). Left panels: amounts were reported by 2h segments. Two-way rANOVA; Wake: treatment, $F(1,14) = 4.128$, $P=0.062$; time: $F(4.253,59.54) = 46.97$, $P<0.0001$; interaction $F(11,154) = 1.751$, $P=0.067$; NREMS, treatment, $F(1,14) = 3.596$, $P=0.079$; time: $F(11,154) = 15.32$, $P<0.0001$; interaction $F(11,154) = 1.452$, $P=0.155$, REMS, treatment, $F(1,14) = 1.578$, $P=0.2296$; time: $F(3.267,45.73) = 22.47$, $P<0.0001$; interaction $F(11,154) = 0.650$, $P=0.7832$. Right panels: amounts were reported by 12h segments. Mann-Whitney tests, **P < 0.01 CTL versus PLX3397. CTL, 7 mice; PLX3397 treated, 9 mice. Data are represented as mean ± SEM.

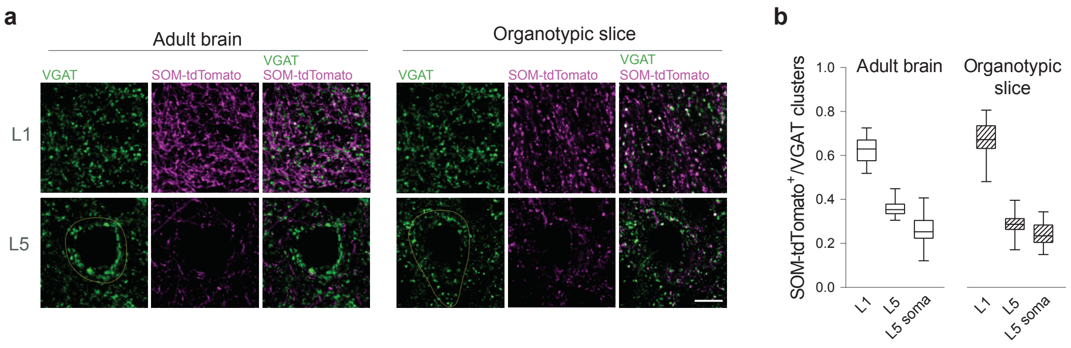


**Figure S3. SOM-IN⁺ inputs are distributed similarly in adult brain and organotypic slices. a**, SOM-IN presynaptic boutons, identified in $SOM^{Cre/+}$:$R26^{tdTom/+}$ adult mice as SOM-tdTomato⁺VGAT⁺ terminals, are denser in upper cortical layers of brain and organotypic slices. Scale bar, 10 μm. **b**, Fraction of VGAT⁺ puncta colocalized to SOM-tdTomato⁺ boutons. Of note, the majority of VGAT⁺ puncta in adult brain and organotypic slices cortical L1 are SOM-IN inputs. Adult brain: n= 31 (L1) and 33 (L5) FOVs from 3 $SOM^{Cre/+}$:$R26^{tdTom/+}$ mice. Organotypic slices: n= 24 (L1), 12 (L5) FOVs from 1 $SOM^{Cre/+}$:$R26^{tdTom/+}$ culture.

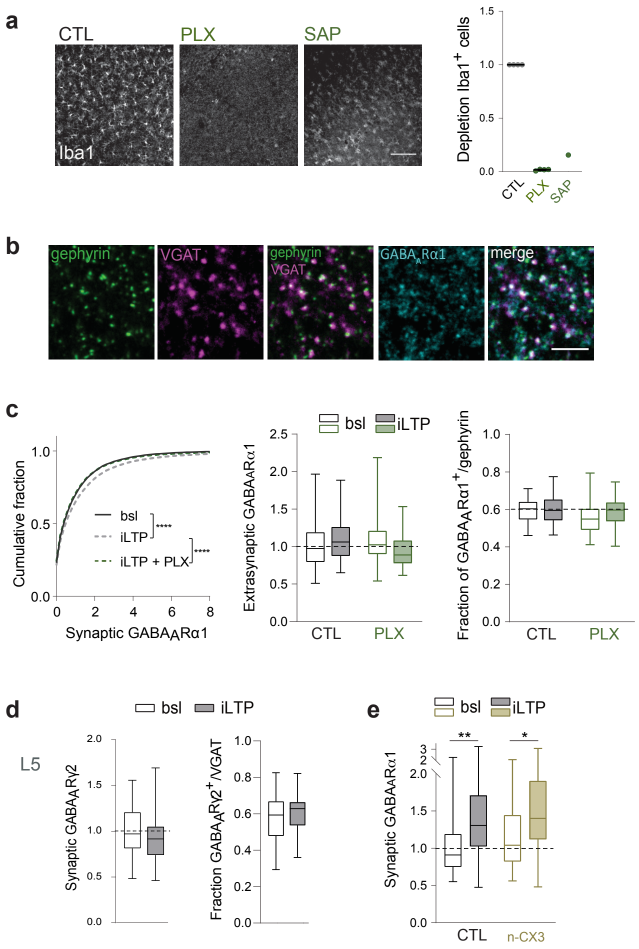


**Figure S4. Microglia are required for iLTP-induced GABA$_A$Rs plasticity. a**, Left: Iba1+ cells are depleted in organotypic slices by PLX3397 (PLX) or Mac1-Saporin (SAP) treatment. Bar, 100 µm. Right: Density of Iba1+ cells normalized to untreated slices (CTL). n= 4 to 1 independent experiments. **b**, Confocal images of inhibitory synaptic markers gephyrin, VGAT and GABA$_A$Rα1 in organotypic slices cortical L1. Scale bar, 5 µm. **c**, Left: Cumulative fraction of the intensity of GABA$_A$Rα1 clusters at gephyrin+ cluster. ****p<0,0001, Kolmogorov-Smirnoff. Middle: Mean intensity of extrasynaptic GABA$_A$Rα1. Right: Fraction of gephyrin+ clusters colocalized to GABA$_A$Rα1. n= 44 to 50 FOVs from 6 independent experiments. **d**, Left: Mean intensity of GABA$_A$Rγ2 clusters at VGAT+ clusters around NeuN+ cell bodies showing no changes in GABA$_A$Rγ2 at putative somatic L5 synapses upon iLTP. Right: Fraction of VGAT+ clusters colocalized to GABA$_A$Rγ2 cluster in putative L5. n= 43 to 45 FOVs from 6 independent experiments. **e**, Mean intensity of GABA$_A$Rα1 clusters at gephyrin+ cluster showing no suppression of iLTP effect by CX3CL1 neutralizing antibody (n-CX3). n= 63 to 69 FOVs from 6 independent experiments. *p<0.05 and **p<0.01, nested one-way ANOVA followed by Sidak's multiple comparison test.

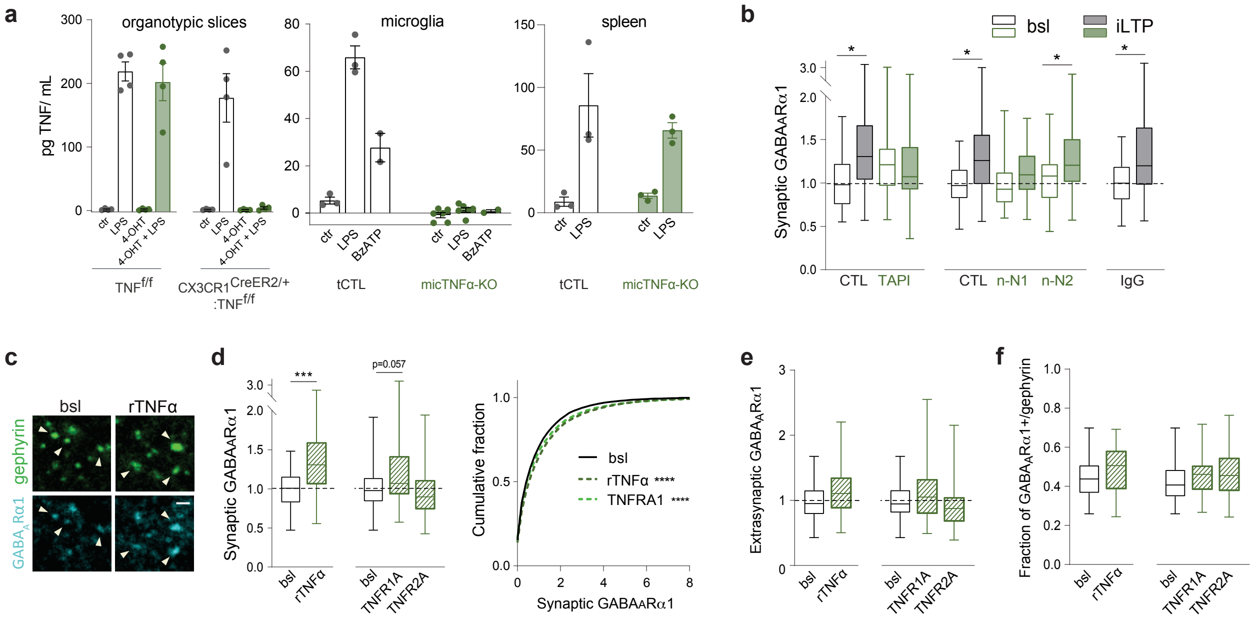


**Figure S5.** **–TNFα controls GABA$_A$R plasticity. a**, Conditional deletion of TNFα from CX3CR1-expressing cells in organotypic slices and *in vivo* assessed by ELISA. Graphs show TNFα concentration (pg/mL) as mean ± SEM. Left: Complete lack of LPS-induced TNFα production by organotypic slices upon 4-OHT-induced recombination of microglial TNFα in a CX3CR1$^{CreERT2/+}$:TNF$^{f/f}$ background but not control slices (TNF$^{f/f}$). n= 2 replicates from 2 individual experiments. Middle: Following feeding with tamoxifen-containing food, LPS- or BzATP-induced TNFα release by adult mouse primary microglia observed in transgenic controls (tCTL, CX3CR1$^{GFP/+}$:TNF$^{f/f}$) but not on microglia-TNFα depleted mice (micTNFα-KO, CX3CR1$^{CreERT2/+}$:TNF$^{f/f}$). n= 2 to 6 mice per group. Right: On spleen cultures from tamoxifen-treated mice, LPS-induced TNFα release is not affected in both tCTL and micTNF-KO, demonstrating microglia-specific TNFα deletion *in vivo*. n= 3 mice per condition. **b**, Mean intensity of GABA$_A$Rα1 clusters at gephyrin$^+$ clusters upon blockade of TNFα cleavage (TAPI), neutralization of TNFR1 (n-N1), TNFR2 (n-N2) and control IgGs (IgG). n= 62 to 89 FOVs from 5-9 independent experiments. **c**, Representative images showing increase of GABA$_A$Rα1 (cyan) at gephyrin$^+$ clusters (arrowhead, green) after 20min treatment with recombinant TNFα (rTNFα). **d**, Effect of rTNFα and TNFR1 activating antibodies (TNFRA1), but not TNFR2 activating antibodies (TNFRA2), on GABA$_A$Rα1 at L1 synapses. Left: Mean intensity of GABA$_A$Rα1 clusters at gephyrin$^+$ clusters. ***p<0.001, nested t-test. Right, Cumulative fraction of the intensity of GABA$_A$Rα1 clusters at gephyrin$^+$ cluster. ****p<0,0001, Kolmogorov-Smirnoff. **e**, Mean intensity of GABA$_A$Rα1 clusters at extrasynaptic sites. **f**, Fraction of gephyrin$^+$ clusters colocalized to GABA$_A$Rα1. **d-f**, n= 65 to 77 FOVs from 8 independent experiments.

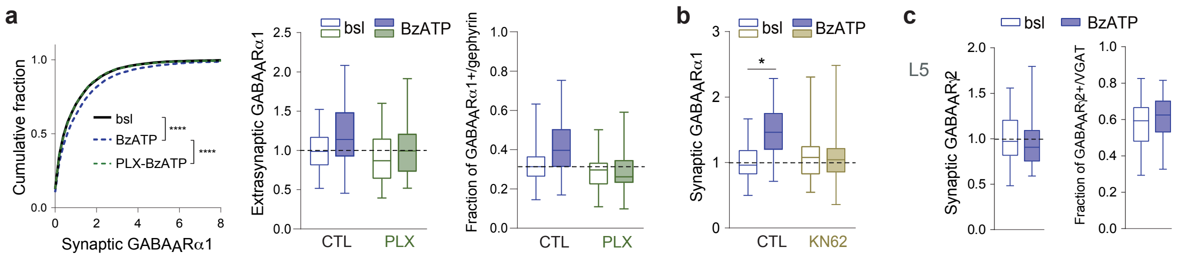


**Figure S6. – BzATP-induced $GABA_AR$ synaptic enrichment. a**, Left: Cumulative fraction of $GABA_AR$α1 clusters' intensities at gephyrin⁺ cluster. ****p<0,0001, Kolmogorov-Smirnoff test. Middle, Mean intensity of extrasynaptic $GABA_AR$α1. Right: Fraction of gephyrin⁺ clusters colocalized to $GABA_AR$α1. n= 49 to 63 FOVs from 5 independent experiments. **b**, Mean intensity of $GABA_AR$α1 clusters at gephyrin⁺ cluster. KN62, a CaMKII inhibitor, abolishes BzATP-induced synaptic upregulation of $GABA_AR$α1, indicative of a shared molecular mechanism at the neuronal level with iLTP. n= 53 to 57 FOVs from 5 independent experiments. *p<0.05, nested one-way ANOVA followed by Sidak's multiple comparison test. **c**, BzATP has no effect on synaptic $GABA_AR$ at putative L5. Left: Mean intensity of $GABA_AR$γ2 clusters at VGAT⁺ clusters. Right: Fraction of VGAT⁺ clusters colocalized to $GABA_AR$γ2 cluster. n= 42 to 45 FOVs from 6 independent experiments.

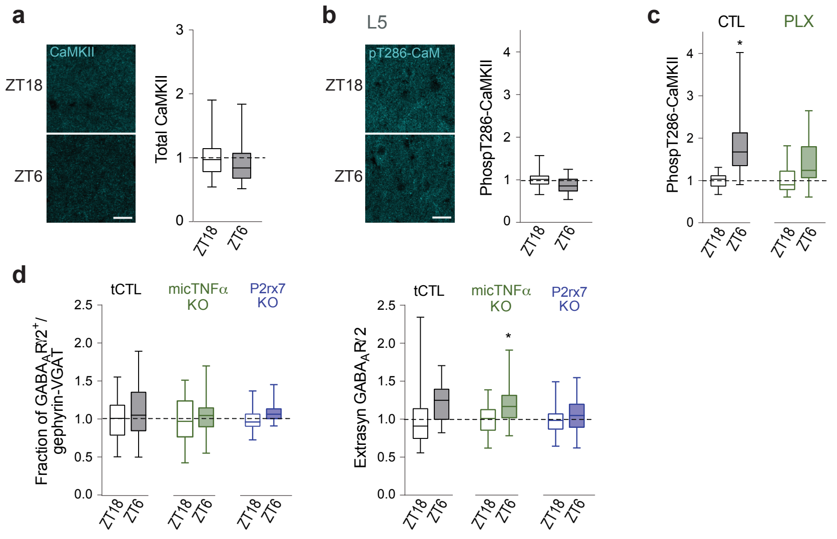


**Figure S7.– Microglia control $GABA_A$R plasticity and fluctuations of CaMKII phosphorylation across the light/dark cycle. a**, Left: Confocal images of L1 total CaMKII immunoreactivity showing no changes between ZT18 and ZT6 in transgenic control mice (tCTL). Scale bars, 20 µm. Right: Mean intensity of total CaMKII signal normalized to ZT18. n= 40 to 50 FOVs from 4-5 mice per group. **b**, Confocal images of L5 Thr286-phosphorylated CaMKII immunoreactivity. Right: Mean intensity of Thr286-phosphorylated CaMKII signal normalized to ZT18. n= 32 to 40 FOVs from 4-5 mice per group. **c**, Mean intensity of Thr286-phosphorylated CaMKII signal in L1 normalized to ZT18. n= 49 to 50 FOVs from 5 mice per group. *p<0.05, nested t-test. **d**, Left: Fraction of gephyrin$^+$VGAT$^+$ synapses colocalized to $GABA_A$R cluster. Right: Mean intensity of $GABA_A$R clusters at extrasynaptic sites. n= 48 to 65 FOVs from 4-5 mice per group. *p<0.05, nested t-test.

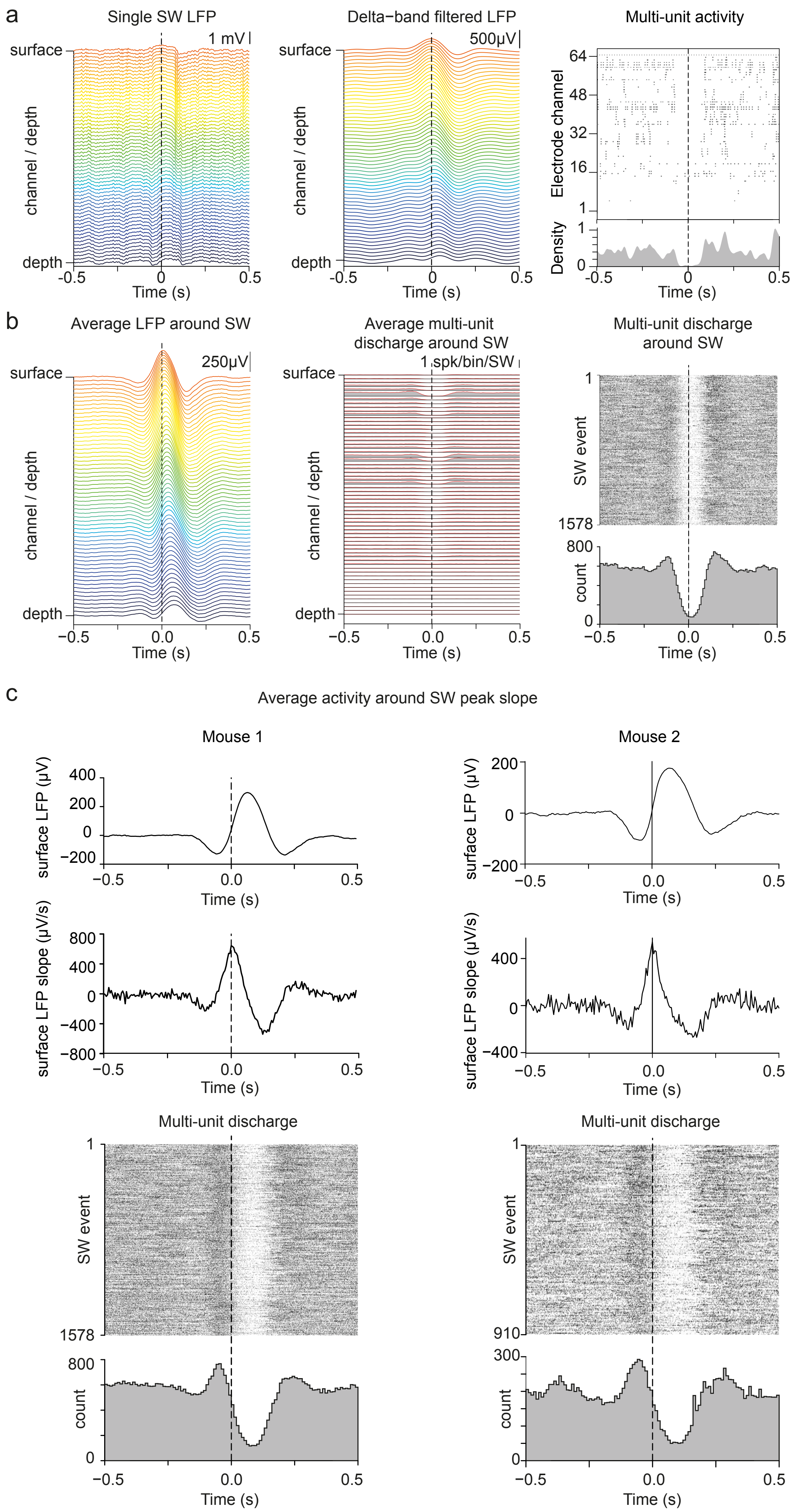

a
Single SW LFP
surface
depth
channel / depth
1 mV
Time (s)
Delta−band filtered LFP
500µV
Multi-unit activity
Electrode channel
Density
b
Average LFP around SW
250µV
Average multi-unit discharge around SW
1 spk/bin/SW
Multi-unit discharge around SW
SW event
count
c
Average activity around SW peak slope
Mouse 1
Mouse 2
surface LFP (µV)
surface LFP slope (µV/s)
Multi-unit discharge

**Figure S8. SWs coincide with cortical down-states and peak upward slope corresponds to the onset of down-states**. **a**, Example of single SW recorded across cortical layers by linear electrodes (with 20μm spacing between the contacts). The channels are plotted and colored by depth. The top channel is located close underneath the surface, and the bottom channel ('depth') is located 1275μm below. Left: raw local field potential (LFP) traces. Middle: delta-band filtered LFP. Right: multi-unit activity across all electrodes channels; each line corresponds to a channel, and each dot represents the time of an extracellular spike detected on the channel. The density of spikes detected on all channels is evaluated using a 10ms Gaussian kernel and plotted at the bottom. **b**, Average LFP and multi-unit activity around the peak of all SW detected in a 2h-recording session. Left: average LFP trace for each depth. Middle: spike count around the peak of the SW; each line corresponds to the cross-correlogram between the times of the peak of SW and the multi-unit activity of a single channel. Right: rasters of the total multi-unit activity detected across all channels for each individual SW (each line corresponds to a SW event, each dot on this line to an extracellular spike); a grand histogram is indicated at the bottom. Note that there is a clear drop of the density of spikes around the peak of each SW indicating the correspondence between the SW and cortical down-states. **c**, The peak upward slope corresponds to the drop of multi-unit cortical activity below the basal firing rate. The results from 2 mice are displayed. Top: average SW waveform at the surface centered on the time of maximal slope of the SW. Middle: average derivative of the surface LFP around the time of maximal slope of the SW. Bottom: raster and histogram of the multi-unit activity for all SWs, centered on the maximal slope of each SW.

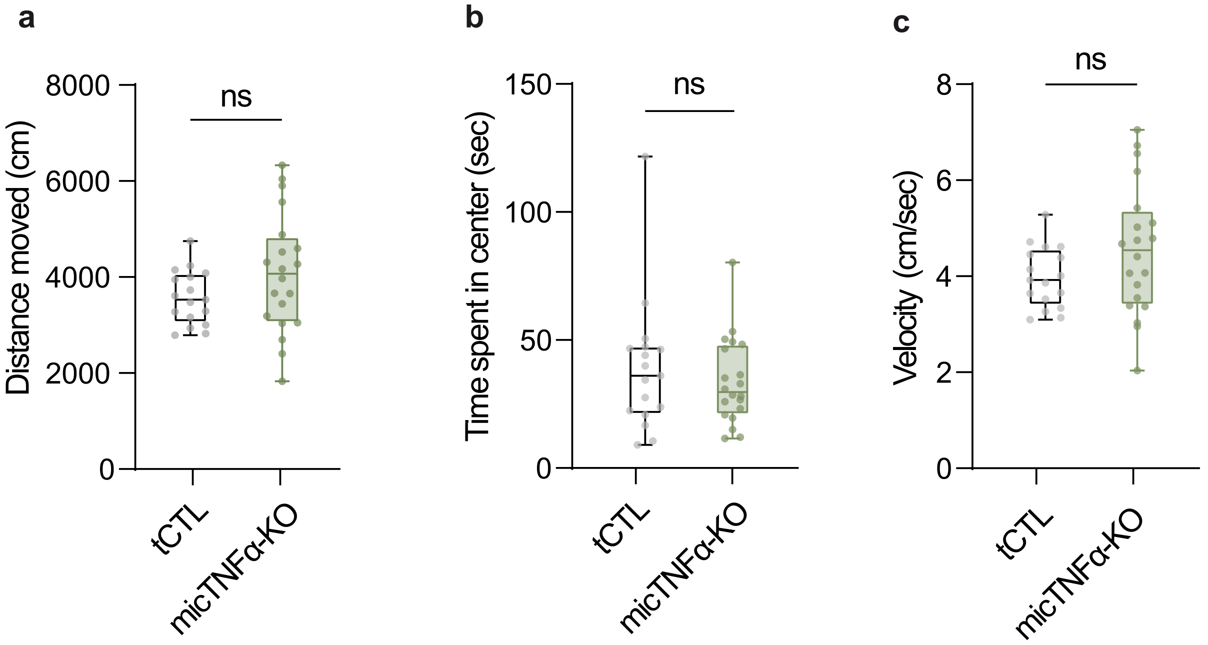


**Figure S9. Microglial TNFα does not modulate locomotor activity and anxiety-like behaviors.** No significant difference were measured between tCTL and micTNFα-KO mice in the open-field test for: **a**, The distance covered (two-tailed unpaired T-test, t(35)=1.533, p=0.1324); **b**, The time spent in the central zone (two-tailed unpaired T-test, t(35)=0.7278, p=0.4716); and **c**, The velocity (two-tailed unpaired T-test, t(35)=1.587, p=0.1214);  (a-c) n=17 tCTL and 20 micTNFα-KO mice. Data are expressed as means ± SEM.

**Extended table 1. Sleep and wake characteristics in control and micTNFα-KO mice.**

| Baseline sleep (24h) | Amounts (min) | | Number of bouts | | Bouts mean duration (min) | |
|---|---|---|---|---|---|---|
| | tCTL | micTNFα-KO | tCTL | micTNFα-KO | tCTL | micTNFα-KO |
| WAKE | 757.7 ± 21.3 | 739.1 ± 16.5 | 209.3 ± 13.5 | 204.9 ± 7.6 | 3.73 ± 0.26 | 3.65 ± 0.18 |
| NREM | 616.7 ± 22.5 | 619.4 ± 16.0 | 205.9 ± 14.0 | 202,5 ± 7,7 | 3.00 ± 0.19 | 3.08 ± 0.11 |
| REM | 65.6 ± 3.0 | 80.4 ± 2.3** | 56.4 ± 3.0 | 76.8 ± 4.2** | 1.17 ± 0.04 | 1.07 ± 0.06 |

The data are means ±S.E.M. Two-tailed unpaired t-test; **P < 0.01, significantly different from transgenic control mice (15 mice per groups).

**Extended table 2. Sleep and wake characteristics in control and PLX3397 treated mice during the light and the dark phase.**

| Baseline sleep (12h - Light) | Number of bouts | | Bouts mean duration (min) | |
|---|---|---|---|---|
| | CTL | PLX | CTL | PLX |
| WAKE | 128 ± 7 | 126 ± 9 | 1.7 ± 0.2 | 1.8 ± 0.1 |
| NREM | 128 ± 7 | 127 ± 9 | 3.6 ± 0.2 | 3.7 ± 0.3 |
| REM | 44 ± 3 | 41 ± 4 | 1.2 ± 0.1 | 1.3 ± 0.1 |

| Baseline sleep (12h - Dark) | Number of bouts | | Bouts mean duration (min) | |
|---|---|---|---|---|
| | CTL | PLX | CTL | PLX |
| WAKE | 68 ± 7 | 76 ± 6 | 8.2 ± 1.0 | 6.2 ± 0.5 |
| NREM | 68 ± 7 | 76 ± 6 | 2.8 ± 0.3 | 3.5 ± 0.3 |
| REM | 11 ± 2 | 13 ± 2 | 1.1 ± 0.1 | 1.1 ± 0.1 |

Data as means ± S.E.M. CTL, 7 mice; PLX3397 treated, 9 mice. Mann-Whitney tests, CTL
